# Establishing toxicologically relevant primary alveolar macrophage models to enable a realistic assessment of nanomaterials safety

**DOI:** 10.64898/2026.10.12.758669

**Authors:** Hongyu Ren, Chenxi Li, Haiyun Zhang, Verena Haefner, Jiating Deng, Inti De la Rosa Velazquez, Jesús Pérez Gil, Carola Voss, Markus Rehberg, Tobias Stoeger, Lianyong Han

**Author notes:** **Contact information** Correspondence to T. Stoeger, L. Han.

## Abstract

Alveolar macrophages (AMs) are key sentinels of the distal lung, maintaining surfactant homeostasis and clearing inhaled particles through continuous surveillance and phagocytosis. Mechanistic studies of particle–cell interactions, important to assess the respiratory safety or toxicity of particles, however, are limited by the lack of suitable *in vitro* models that preserve the core features of resident AMs. Here, we benchmarked six murine AM models, including short-term cultured primary AMs (AMs 3 h, AMs 24 h), primary AMs maintained with GM-CSF, TGF-β, and the PPAR-γ agonist rosiglitazone (AMs GTR), bone marrow-derived AM-like cells (BM-AMs), the immortalized AM cell line MH-S, and the non-transformed fetal liver-derived AM-like cell line MPI, against *in vivo* AMs based on surface phenotype, transcriptomic identity, and their responses to three carbon-based nanomaterials (CBNs). Both cell lines showed limited AM identity, characterized by low CD11c and SiglecF but high CD11b surface expression. In contrast, AMs GTR and BM-AMs more closely preserved resident AM-like surface marker profiles. Transcriptomic features, although differing for alterations in metabolism-related pathways, particularly oxidative phosphorylation, revealed most *in vivo-*like for the primary AM models. Functional, toxicological assessment using soot-like carbon nanoparticles (CNP), double-walled carbon nanotubes (DWCNT) and multi-walled carbon nanotubes (MWCNT) demonstrated strong model-dependent responses. Consistent with MWCNT-induced reduction in AM numbers observed *in vivo*, AMs GTR and BM-AMs exhibited the strongest *in vitro* responses, including cytotoxicity and inflammatory cytokine release. MH-S cells in contrast stood out for their cytotoxic sensitivity supporting their use for high throughput cytotoxicity tests. Overall, these findings identify AMs GTR and BM-AMs as the closest models for studying AMs biology and AM-intrinsic responses to poorly soluble nanoparticles like CBNs, and with regard to availability of higher cell numbers, BM-AMs might provide a scalable platform for mechanistic and higher-throughput nanotoxicology studies.

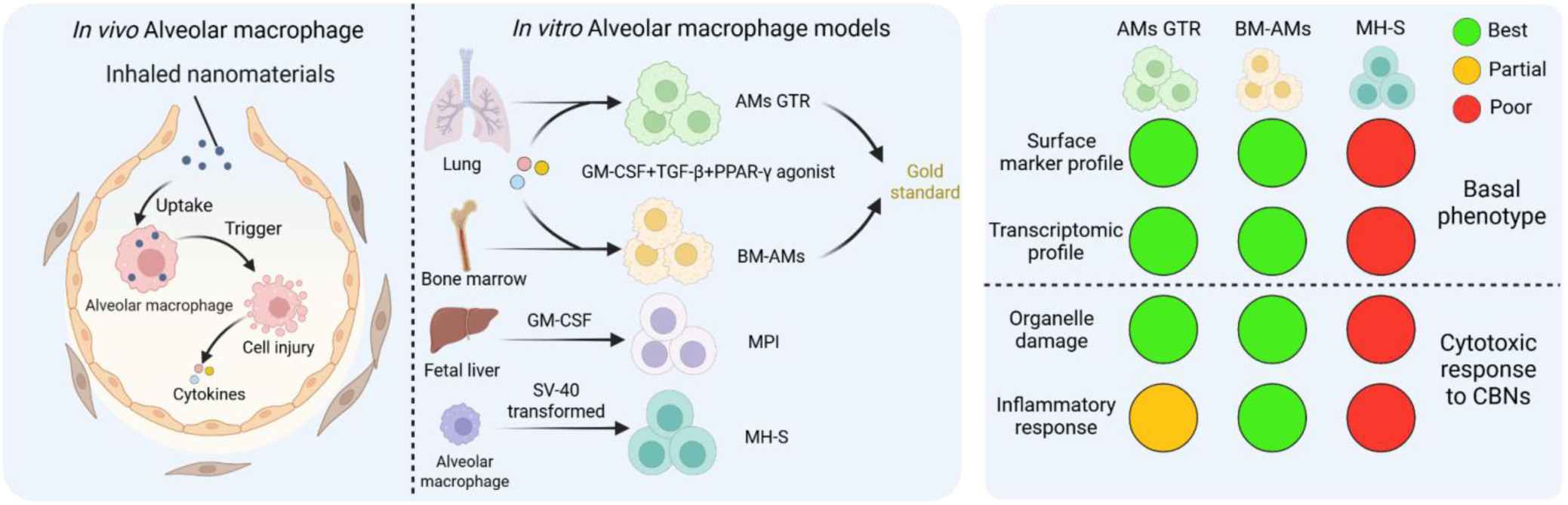

## Introduction

With the inevitable emission of combustion-derived carbon nanoparticles (CNPs) and the increasing use of engineered CNPs and carbon nanotubes (CNTs) in industry and nanomedicine, concerns have been raised about the release of these carbon-based nanomaterials (CBNs) into the environment and in particular their potential risks to human health, especially via inhalation.^1,2^ In addition, engineered CBNs used in industrial applications, particularly carbon nanotubes, may pose respiratory hazards following occupational or environmental exposure, emphasizing the need for reliable and practicable approaches to their hazard and risk assessment.^3^ Building on approaches initially developed for cosmetic ingredients,^4^ Next Generation Risk Assessment (NGRA) has been defined as an exposure-driven framework that integrates multiple lines of evidence using New Approach Methodologies (NAMs), with the aim of assessing risk without the use of vertebrate animal studies. This world-wide call has boosted the search for reliable *in vitro* models tremendously but critical studies also disclosed discrepancies between risk and hazard data derived from *in vivo* or *in vitro*. For respiratory toxicity testing, the so-called poorly soluble particles of low toxicity, revealed to be tricky when tested in cell lines at the *in vitro* level. While in the lungs of animals, all materials seem to induce inflammatory response levels, depending on the particle surface area deposited in the lung,^5^ this correlation revealed to be difficult being reproduced at the *in vitro* level.^6^ In addition to surface area, particle shape is an important determinant of poorly soluble particles. CNPs, the particulate basis of soot, are spherical shaped nanoparticles consisting of primary particles (10-100 nm), forming aggregates (85-500 nm) and agglomerates (1-100 μm). This hierarchical structure grants CNPs a high surface area to volume ratio,^7^ which enables pro-oxidative particle-cell interactions thereby contributing to acute and transient lung inflammation.^8^ Depending on the number of layers of contained graphene sheets, CNTs can be classified into single-walled (SWCNT), double-walled (DWCNT), and multi-walled carbon nanotubes (MWCNT).^9^ Due to their elongated, fiber-like shape and biopersistent properties, high doses or continuous exposure of CNTs are well-known to induce chronic lung diseases (CLDs) in mouse studies, like lung fibrosis, asthma, chronic obstructive pulmonary disease (COPD) and even lung cancer,^10,11^ as exemplified by asbestos-like MWCNT.

A closer look into the cellular anatomy of lungs is required when striving for alternative lung cell model systems for toxicology testing strategies.^12^ As the first line of defense in the distal lung, alveolar macrophages (AMs), self-maintained by controlled proliferation, constantly patrol the alveolar space to clear inhaled pathogens and foreign materials to maintain lung homeostasis.^13–15^ After deposition in the alveolar space, particles such as CBNs are recognized and internalized by AMs through opsonin-assisted, scavenger-receptor-mediated and complement pathways and are subsequently delivered to phago- lysosomal compartments for degradation or sequestration.^16–18^ The internalization and intracellular processing of CBNs, particularly fibers, can be accompanied by AM activation or damage, thereby contributing to the initiation of acute lung inflammation.^19,20^ The physicochemical properties of CBNs are key determinants of the extent, type and durability of AM damage, thereby influencing whether the initial inflammatory response resolves or progresses to prolonged tissue damage.^21^ For example, spherical CNP typically elicit robust acute neutrophilic inflammation and oxidative stress, whereas their effects on AM state and numbers appear moderate and may restore over days to weeks depending on particle burden and dispersion.^7,22^ DWCNT likewise exhibit biopersistence and can drive chronic inflammation and fibrotic and even neoplastic pathologies in multi-year studies, supporting a fiber-hazard paradigm.^23,24^ Exposure to long, stiff MWCNT can induce frustrated phagocytosis in AMs and mechanically disrupt lysosomal integrity, triggering cathepsin release, mitochondrial stress and ultimately inflammatory cell death.^25,26^ A repeated AM activation and injury may further compromise homeostasis, and amplify the inflammatory cascade, thereby promoting CLDs.^27,28^

Given the central role of AMs in linking CBNs toxicity to lung pathogenesis, we have previously shown that short- and long-term exposure of mice to needle like, fiber-shaped MWCNT induced a significant reduction in the CD11c^+^, F4/80^+^ and CD11b^-^ cell population - representing a canonical cell surface marker profile of resident AMs. This AM depletion persisted throughout the 28-day observation period, suggesting insufficient replenishment of resident AMs following MWCNT-induced cell loss.^20,29^ A similar AM depletion effect was observed in DWCNT exposed mice, however, less prominent, while CNPs were least toxic to AMs.^20^ Mechanistically, the detrimental effects of CBNs on the fate of AMs has been attributed to multiple factors, including the formation of a protein corona on the surface of CBNs aggregates in the surfactant-rich alveolar environment, triggering cell circuits between AMs, epithelial cells and recruited inflammatory cells.^30,31^ To date animal studies remain essential for evaluating the full toxicity of inhaled particles such as CBNs. While there is an urgent need for suitable *in vitro* models, recapitulating their mode of action in defined cell culture systems, the developments of such new approach methodologies (NAM) remain especially difficult to reach for poorly soluble particles such as CBNs.^6,32^ Therefore, it is essential to establish *in vitro* AM models that reliably recapitulate *in vivo* AM identity and function, enabling controlled investigation of AM-intrinsic responses to particle exposure.

It is well known that core features of AM, including their anti-inflammatory polarization state, their specific lipid metabolism and oxidative phosphorylation program, as well as their patrolling behavior, are largely shaped by the alveolar niche.^13^ Accordingly several attempts have been made to establish different AM models that recapitulate their core functions. For example, primary AMs, isolated from mice were transformed into an immortalized cell line (MH-S) using a lentivirus SV40 system.^33^ Non-transformed AM-like cells (MPI) were generated from murine fetal liver monocytes during culture with GM-CSF and can be expanded *in vitro* for at least 90 passages without apparent morphological or functional changes.^34^ For rats, the alveolar macrophage cell line (NR8383) was generated by extended over 36 month culture to finally subcloning self-proliferating clones.^35^ While this cell line is effectively used for toxicological investigations, no comparable tool is available for mice.^36^ More recently, a cytokine cocktail consisting of GM-CSF, TGFβ and a PPARγ agonist (GTR) was shown to differentiate murine bone marrow derived monocytes into AM-like cells (BM-AMs) expressing high levels of the AM prototypic surface markers CD11c and SiglecF.^37^ However, it remains unclear to what extent these *in vitro* AM models recapitulate resident AM identity, including niche-imprinted transcriptional programs and functional behaviors relevant to CBNs exposure.

By comparing the described murine *in vitro* AM models to *in vivo* AMs from mice for their cellular phenotype, their transcriptomic landscape, inflammatory cytokine release profile and intracellular organelle response towards distinct CBNs, we provide evidence that (i) both primary AM models, AMs GTR and BM-AMs retain the *in vivo*-like AM phenotype best; (ii) both AMs GTR and BM-AMs retain several key transcriptional pathways, even if however, deficient in oxidative phosphorylation; (iii) BM-AMs share a similar inflammatory cytokine release profile with that detected in vivo in bronchial lavage fluid from MWCNT exposed mice;^20^ and finally (iv) both BM-AMs and AMs GTR show comparable intracellular organelle damage pattern in response to MWCNT. Along with the ability to proliferate and thus producing cell numbers at high-yield, BM-AMs might therefore serve as a promising murine *in vitro* AM model to be implemented in more complex cell systems and mechanistic studies of respiratory toxicology or biomedicine.

## Results

### Phenotypic comparison of different *in vitro* models of murine alveolar macrophage

To systematically evaluate the suitability of murine *in vitro* AM models for respiratory particle toxicity assessment, we established and characterized six distinct cell models and compared them to freshly isolated AMs by bronchoalveolar lavage (BAL) from young and healthy mice as an *in vivo* reference. The models included: (1) freshly isolated AMs cultured for 3 h (AMs 3 h); (2) freshly isolated AMs cultured for 24 h (AMs 24 h); (3) primary AMs cultured for 7 days with a GTR-cocktail consisting of GM-CSF, TGF-β, and the PPARγ-agonist rosiglitazone (AMs GTR); (4) bone marrow-derived AM-like cells cultured with GTR (BM-AMs); (5) the SV40-transformed MH-S cell line; and (6) the non-transformed MPI cell line.

As expected, freshly isolated BAL-AMs were predominantly positive for the two prototypic AM markers CD11c⁺ (91.0 ± 0.8%) and SiglecF⁺ (94.6 ± 1.5%), whereas only 2.2 ± 0.3% of cells were CD11b⁺ positive. The latter together with CD18 forms Macrophage-1 antigen (Mac-1) and serves as a common marker for macrophages and other bone marrow derived leukocytes. After 3 h of culture, the percentage of CD11c⁺ cells decreased markedly to 22.9 ± 3.3%, accompanied by a more moderate reduction in SiglecF⁺ cells to 77.3 ± 5.9%, while CD11b⁺ cells remained scarce (4.6 ± 1.4%). These changes indicate a rapid, culture- associated loss of the typical AM surface phenotype. By 24 h, the percentage of CD11c⁺ cells partially recovered to 63.8 ± 1.2%, suggesting that the early phenotypic changes were at least partly reversible and illustrating the plasticity of AM surface marker profiles during *in vitro* culture (**Fig. 1A, B**). AMs GTR closely recapitulated the surface phenotype of resident AMs, with high percentages of CD11c⁺ (79.9 ± 9.1%) and SiglecF⁺ cells (98.3 ± 0.8%), although a moderate fraction remained CD11b⁺ (20.2 ± 7.0%). BM-AMs were also predominantly CD11c⁺ (91.2 ± 1.7%) but showed an intermediate proportion of SiglecF⁺ cells (52.2 ± 2.7%) and a high percentage of CD11b⁺ cells (82.8 ± 4.5%), consistent with their bone marrow-derived origin (**Fig. 1A, B**). In contrast, MH-S and MPI cells showed little or no positivity for the canonical AM markers CD11c and SiglecF, while remaining predominantly CD11b⁺. Together, these results reveal a spectrum of AM-like phenotypes across the investigated models and demonstrate substantial plasticity of AM surface marker profiles under different *in vitro* culture conditions (**Fig. 1A, B**).

**Fig. 1.**
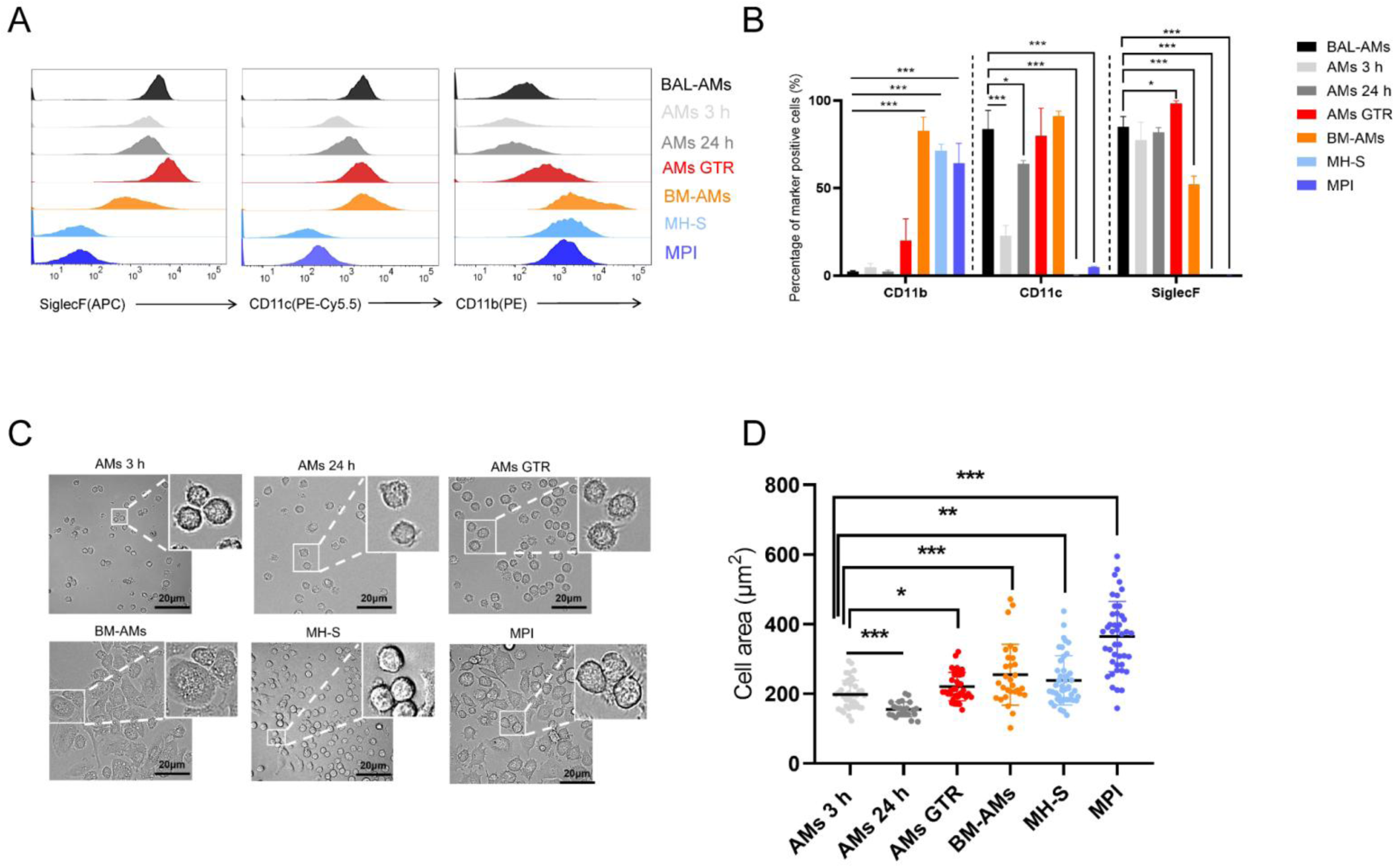
AMs GTR and BM-AMs recapitulate major phenotypic features of resident alveolar macrophages. **(A)** Representative flow cytometry histograms showing the expression profiles of alveolar macrophage identity markers (SiglecF and CD11c) and the monocyte-associated marker CD11b across freshly isolated BAL-AMs and different *in vitro* AM models. **(B)** Quantification of flow cytometric analysis of CD11b+, CD11c+, SiglecF+ in the AM models (n = 3). Values are shown as mean ± SEM. Data were analyzed by using one-way ANOVA followed by Tukey’s multiple comparisons test. “*” indicates statistically significant difference versus “BAL-AMs”. \**P* < 0.05, \*\**P* < 0.01, \*\*\**P* < 0.001. **(C)** Representative bright-field images showing morphological differences among *in vitro* AM models, with increased spreading observed in AMs GTR and BM-AMs. Scale bar = 20 μm. **(D)** Quantification of projected cell area from individual cells of different AM models. A total of 24-43 individual cells per model were analyzed from three independent experiments. Data are presented as mean ± SEM. Statistical significance was determined using one-way ANOVA followed by Tukey’s multiple comparisons test. “*” indicates statistically significant difference versus “AMs 3 h”. \**P* < 0.05, \*\**P* < 0.01, \*\*\**P* < 0.001.

Comparing the morphological characteristics of these in vitro cultured models (**Fig. 1C, D**) AMs 24 h exhibited the smallest cell size, followed by AMs 3 h. Short-term cultured AMs displayed a predominantly spherical shape, whereas AMs GTR displayed spread morphology with extended pseudopods, indicating strong attachment and related cytoskeletal organization. BM-AMs showed the most pronounced spreading, adopting a characteristic “fried egg” morphology previously described for adherent alveolar macrophages maintained under *in vitro* culture conditions.^38^ MH-S cells were rounded and semi-adherent, whereas MPI cells showed a heterogeneous mixture of spread-out and round morphologies (**Fig. 1C**).

### AMs GTR and BM-AMs retain key transcriptomic features of resident alveolar macrophages

We next evaluated the transcriptomic similarity between the *in vitro* AM models and lung-resident AMs by comparing the bulk RNA-seq datasets generated from the 6 culture models described in **Table 1** with scRNA-seq-derived expression profiles of resident AMs.^20^ To minimize platform-related differences and enable cross-dataset comparison, 13 candidate housekeeping genes were initially evaluated across the bulk and scRNA-seq datasets. *Hprt* and *Gapdh* showed robust and reproducible expression across both datasets and were therefore selected as reference genes for normalization (**Supplementary Fig. 1**). Principal component analysis (PCA) revealed three distinct clusters (**Fig. 2A**). The AM-like cell lines MH- S and MPI clustered together and were clearly separated from the four primary AM models (pAMs), with the resident scRNA-seq AMs as the most distinct cluster. To further assess the preservation of AM identity across the *in vitro* models, a core set of 483 AM signature genes were selected (**Supplementary Table S1**). The heatmaps illustrate substantial differences in the expression patterns of these genes across models (**Fig. 2B**). Nevertheless, Pearson correlation analysis showed that all primary AM models retained relatively high similarity to resident AMs for the AM signature gene set (Pearson’s *r* > 0.70), whereas MH- S and MPI showed markedly lower concordance (*r* < 0.55) (**Fig. 2C**). A similar trend was observed when the analysis was extended to the whole transcriptome (**Fig. 2D**). Consistent with these findings, a considerable proportion of AM signature genes showed reduced expression in both cell lines (**Fig. 2B**).

**Fig. 2.**
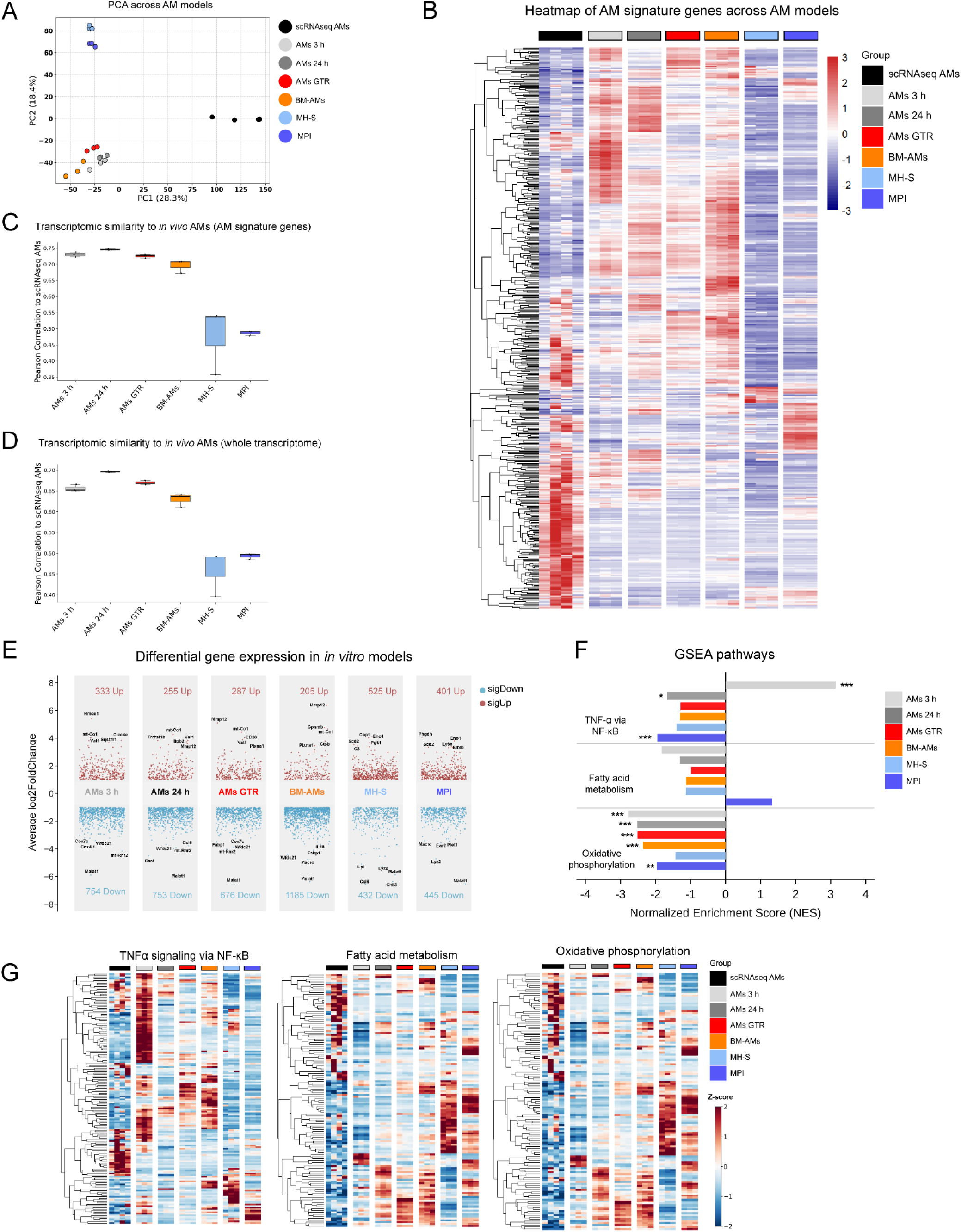
Transcriptomic characterization of established *in vitro* alveolar macrophage models. Bulk RNA sequencing was performed to compare the transcriptomic profile between the 6 distinct *in vitro* models and scRNA-seq data. **(A)** Principal component analysis (PCA) showing the global transcriptomic relationship between the different *in vitro* AM models and resident AMs. **(B)** Heatmap showing the expression patterns of 483 core AM signature genes across resident AMs and the established *in vitro* AM models. Expression values are displayed as row-wise Z-scores. **(C, D)** Pearson correlation analysis comparing the transcriptomic similarity of each *in vitro* AM model with resident AMs based on the 483 AM signature genes (**C**) and the whole transcriptome (**D**). **(E)** Volcano plots showing differentially expressed genes in each *in vitro* AM model relative to resident AMs. Differentially expressed genes were defined by |log2FC| > 1 and FDR < 0.05. Numbers indicate the total numbers of significantly upregulated and downregulated genes in each comparison. **(F)** Gene set enrichment analysis (GSEA) showing normalized enrichment scores (NES) for selected Hallmark pathways, including TNF-α signaling via NF-κB, fatty acid metabolism, and oxidative phosphorylation, across the investigated *in vitro* AM models relative to resident AMs. **(G)** Heatmaps illustrate the expression patterns of genes belonging to TNF-α signaling via NF-κB, fatty acid metabolism, and oxidative phosphorylation pathways. GSEA was performed in R (v4.5.0) using the clusterProfiler package (v4.16.0). Resident AM scRNA-seq data were obtained from GEO accession GSE273313.

**Table 1.** Establishment of different alveolar macrophage *in vitro* models.

| *In vitro* models | Origin | Culture conditions |
| --- | --- | --- |
| BAL-AMs | BAL | Freshly isolated from BAL |
| AMs 3 h | BAL | Cultured for 3 h without GTR in low glucose DMEM |
| AMs 24 h | BAL | Cultured for 24 h without GTR in low glucose DMEM |
| AMs GTR | BAL | Cultured with GTR cocktail for 7 d in low glucose DMEM |
| BM-AMs | Bone marrow | Cultured with GTR cocktail in low glucose DMEM |
| MH-S | SV40 transformed BAL-AMs | Cultured in RPMI 1640 |
| MPI | Fetal liver | Cultured in RPMI 1640 supplied with GM-CSF |

(GTR cocktail: GM-CSF, TGF-β , and peroxisome proliferator-activated receptor $\gamma$ (PPAR-γ ) agonist rosiglitazone)

**Table 2.**
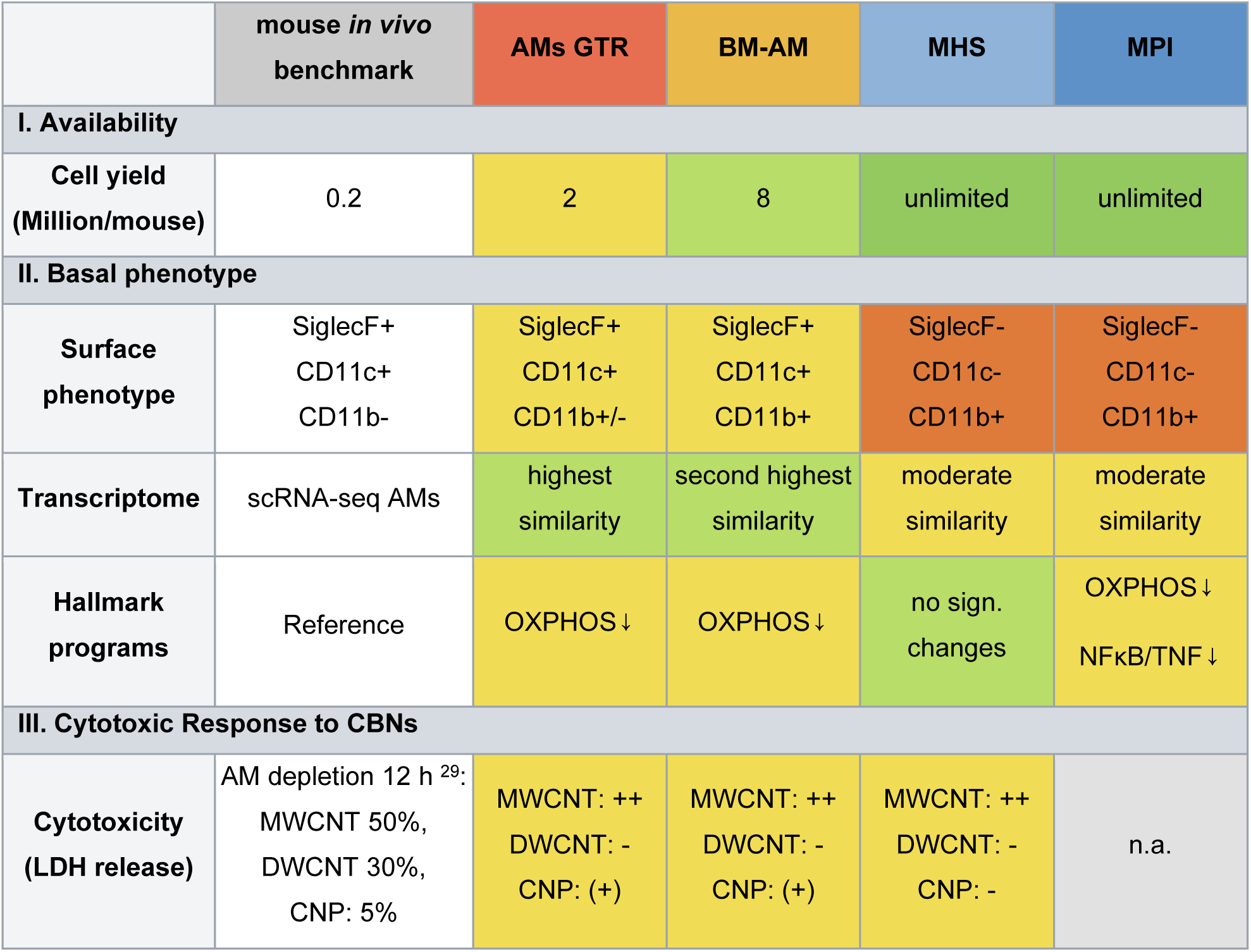
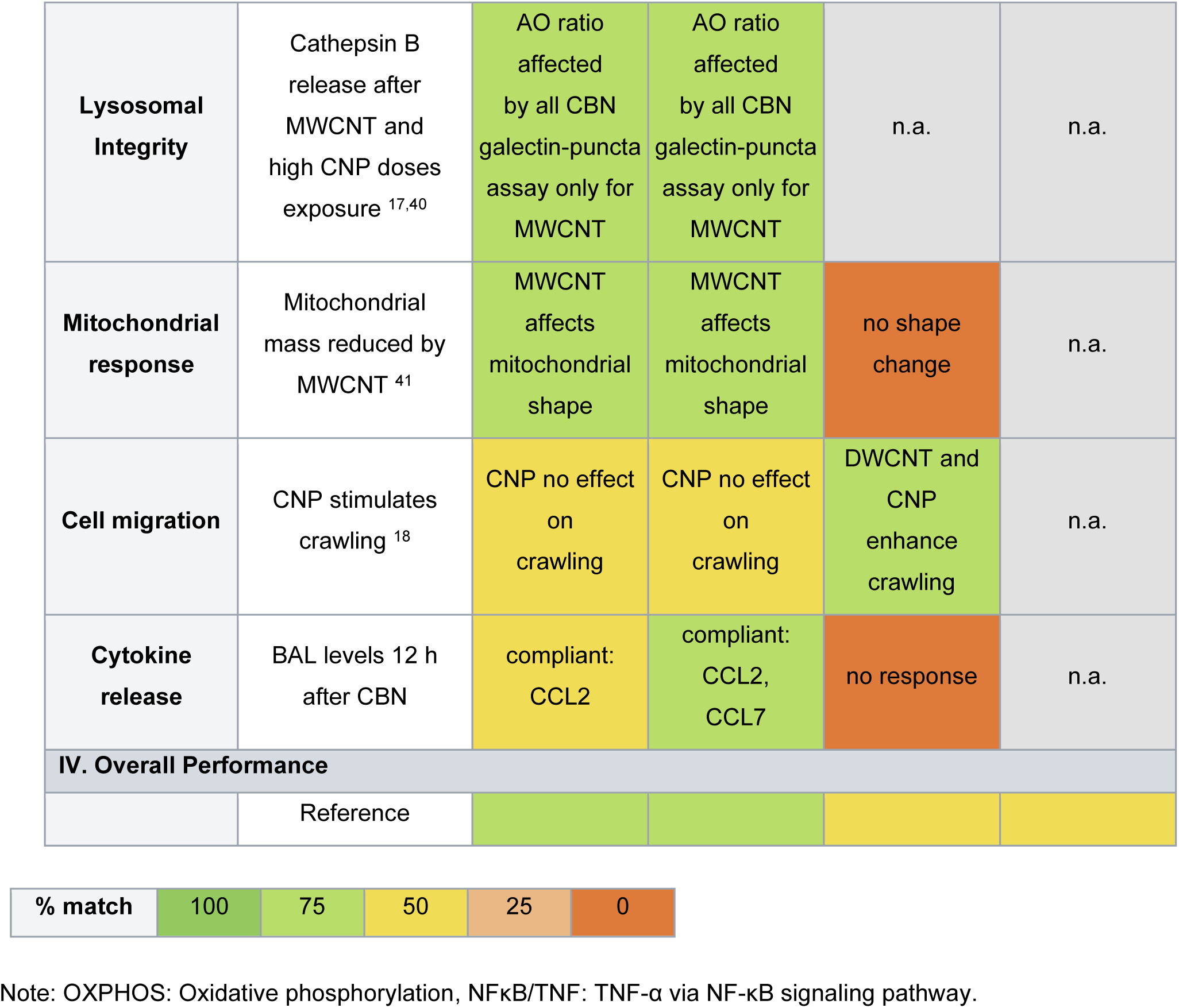
Benchmarking *in vitro* AM models against *in vivo* AMs.

Differentially expressed gene (DEG) analysis and gene set enrichment analysis (GSEA) were further performed. Compared with resident AMs, all established *in vitro* models exhibited extensive transcriptional reprogramming, with the genes *Mt-Co1* (mitochondrial gene), *Vat1*, and *Mmp12* up and *Mt-Rnr2*, *Wfdc21*, and *Malat1* down regulated in the primary cell models and *Eno1*, *Sod2*, up and *Lyz2* and *Malat1* down in the cell line models (**Fig. 2E**). To determine the downstream consequences of these transcriptional differences, pathway enrichment analysis was subsequently performed. TNF-α signaling via NF-κB was selectively enriched only in AMs 3 h, suggesting a transient inflammatory activation associated early with the post-isolation state that diminished after prolonged culture (**Fig. 2F**). In contrast, oxidative phosphorylation was consistently downregulated across all the primary AM models relative to resident AMs, indicating a broad metabolic shift under *in vitro* conditions. Fatty acid metabolism was also reduced in AMs 3 h, whereas this effect was less pronounced in the longer-term primary AM models (**Fig. 2F**). Hallmark “glycolysis” showed modest enrichment in BM-AMs (NES = 1.22, FDR *q* = 0.172) and more pronounced enrichment in AMs GTR (NES = 1.56, FDR *q* = 0.018) relative to resident AMs (**Supplementary Fig. 6**). Thus, despite the consistent alterations in oxidative phosphorylation and fatty acid metabolism, only limited enrichment of glycolytic programs was observed in these two long-term primary AM models. Transcriptional alterations in MH-S and MPI were predominantly associated with proliferative programs, including ribosomal/translation-related processes and Myc–E2F/cell-cycle signaling, together with reduced expression of macrophage-associated signatures (**Supplementary Fig. 2**).

Overall, these analyses indicate that AMs GTR and BM-AMs retain a greater degree of resident AM-like phenotypic and transcriptional identity than the investigated cell lines, while still displaying substantial *in vitro* transcriptional adaptations. Nevertheless, their comparatively preserved AM characteristics support their use as biologically relevant *in vitro* models for further mechanistic studies in response to inhaled nanomaterials.

### Model-dependent cytotoxic responses to carbon-based nanomaterials

Based on the phenotypic and transcriptomic characterization, AMs GTR and BM-AMs were selected as representative *in vitro* models for subsequent functional studies, while MH-S cells were included as a commonly used immortalized alveolar macrophage cell line. To examine whether these models reproduced the material-dependent AM depletion pattern previously observed in the lungs of mice *in vivo* (**Fig. 3A**),^29^ cells were exposed to the same three carbon-based nanomaterials (CBNs), dispersed in whole lung surfactant (physicochemical characteristics are summarized in **Supplementary Table S2 and Fig. S3**), and cell viability and plasma membrane integrity were assessed after 24 h using WST-1 and LDH release assays, respectively.

**Fig. 3.**
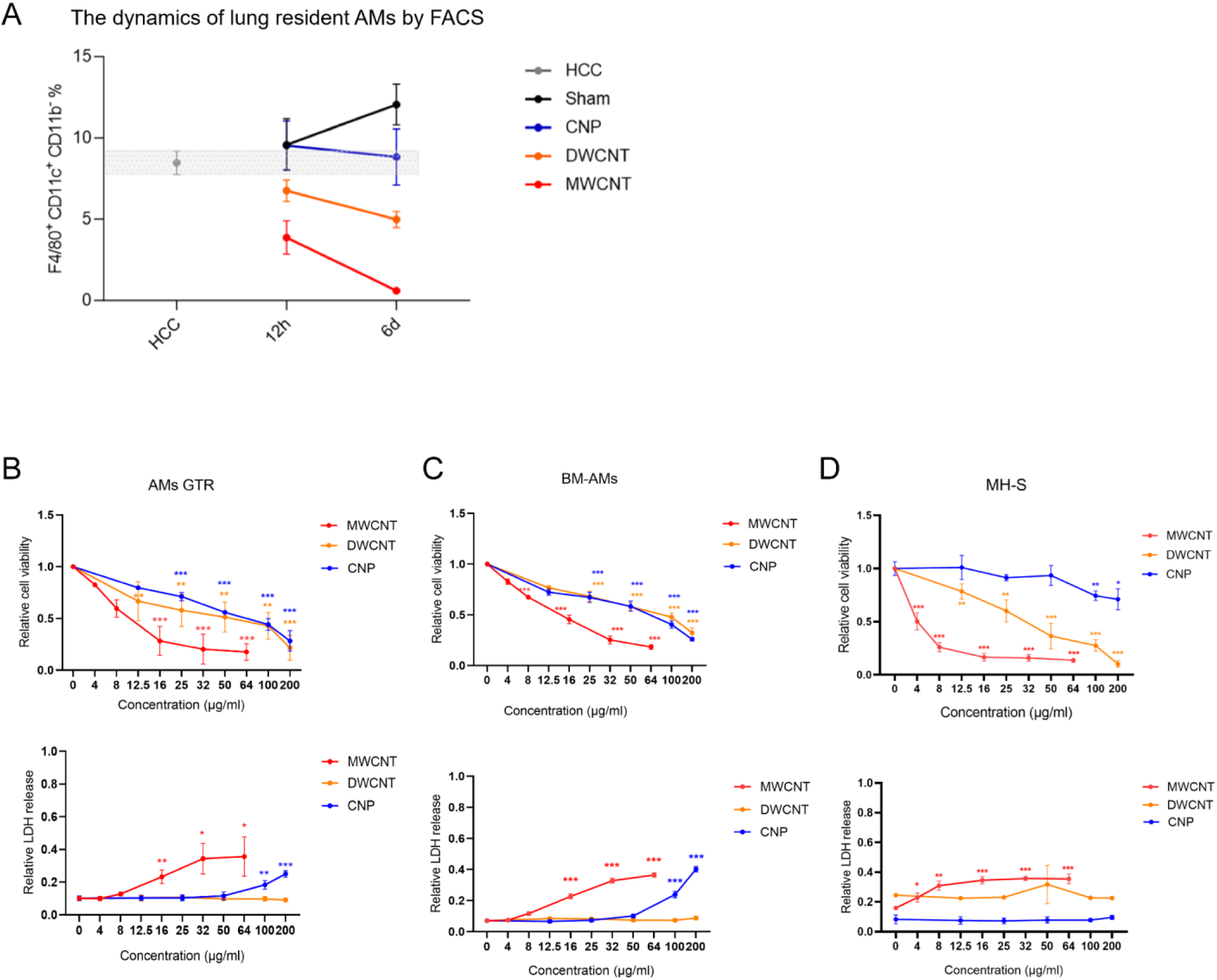
*In vivo* and *in vitro* alveolar macrophage responses to carbon-based nanomaterials. **(A)** Dynamics of lung-resident alveolar macrophages following exposure to carbon nanoparticles (CNP), double- walled carbon nanotubes (DWCNT), or multi-walled carbon nanotubes (MWCNT). Mice (n = 4) were exposed to CNP (20 μg; 54 cm^2^/lung), DWCNT (50 μg; 330 cm^2^/lung), or MWCNT (15 μg; 3.9 cm^2^/lung), and lung-resident AMs were quantified by flow cytometry at 12 h and 6 d. **(B–D)** Cytotoxic responses of AMs GTR (**B**), BM-AMs (**C**), and MH-S cells (**D**) following 24 h exposure to MWCNT (4, 8, 16, 32, and 64 μg/ml), DWCNT (12.5, 25, 50, 100, and 200 μg/ml), or CNP (12.5, 25, 50, 100, and 200 μg/ml). Cell viability was assessed using the WST-1 assay, and plasma membrane damage was evaluated by LDH release. Data are presented as mean ± SEM from three independent experiments (n = 3). \**P* < 0.05, \*\**P* < 0.01, \*\*\**P* < 0.001.

CBNs exposure reduced cell viability in a dose-dependent manner, but the response profiles differed markedly between the primary AM models and MH-S cells (**Fig. 3B–D**). MH-S cells were particularly sensitive to MWCNT exposure, with an approximate IC50 of 4 μg/ml, whereas AMs GTR and BM-AMs showed IC50 values of approximately 13-16 μg/ml. In contrast, MH-S cells were comparatively resistant to CNP exposure. LDH release largely paralleled the viability results, although distinct model-dependent patterns were observed. Following MWCNT exposure, MH-S cells displayed a rapid increase in LDH release at relatively low concentrations (4-16 μg/ml), followed by a plateau at higher concentrations. In contrast, AMs GTR and BM-AMs exhibited a more gradual, concentration-dependent increase in LDH release, reaching approximately 3.5- to 5.1-fold above control levels at 64 μg/ml. DWCNT exposure did not induce a significant increase in LDH release in any of the three models. Notably, CNP exposure increased LDH release only in the primary AM models at the highest concentrations tested, whereas MH- S cells showed little or no membrane damage under the same conditions.

Taken together, these findings demonstrate that cytotoxic responses to CBNs are both material- and model-dependent. The immortalized MH-S cell line displayed a distinct susceptibility profile compared with the established primary AM models, particularly in response to MWCNT and CNP exposure. These differences emphasize the importance of model selection when investigating alveolar macrophage injury mechanisms in nanotoxicology.

### Model-dependent inflammatory cytokine responses to carbon-based nanomaterials

AMs GTR and BM-AMs retained key phenotypic and transcriptomic features of resident alveolar macrophages, although both models exhibited substantial adaptation to *in vitro* culture conditions. Moreover, both models displayed broadly similar patterns of MWCNT-induced cellular injury, with reduced metabolic activity and increased membrane damage, whereas DWCNT and CNP generally produced weaker effects. We therefore next investigated whether these material-dependent injury patterns were accompanied by distinct inflammatory cytokine responses. For this purpose, the AM-models were exposed to CBNs doses causing significant but not excessive effects on cell viability (MWCNT: 16-64 μg/ml, and DWCNT and CNP: 50-200 μg/ml).

MWCNT exposure elicited the strongest inflammatory response in both AMs GTR and BM-AMs, whereas DWCNT and CNP induced much weaker and more selective cytokine changes (**Fig. 4A**). Despite this overall similarity, the magnitude and the number of cytokines induced differed between the two primary AM models. AMs GTR displayed a relatively restricted inflammatory profile following MWCNT exposure, whereas BM-AMs exhibited a broader response involving multiple pro-inflammatory cytokines and chemokines, including prominent induction of TNF-α, IL-6, and IL-1β. In contrast, DWCNT showed a moderate cytokine response only, mostly limited to CCL2 and -7 restricted to BM-AMs, and similar for CNP with a broader response (IL-2, -6, and CCL2, -5, and -7) but limited intensity profile.

**Fig. 4.**
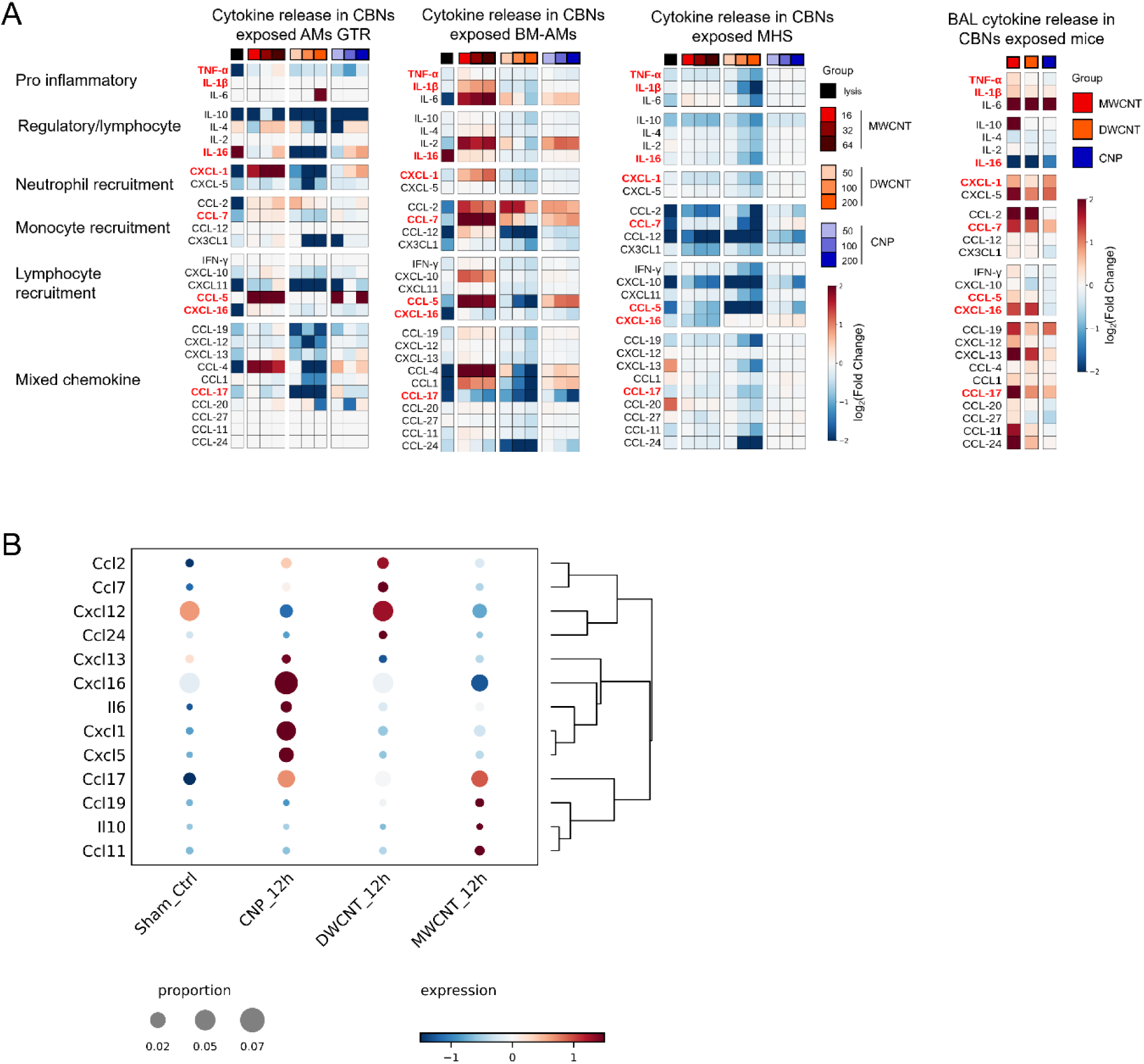
Inflammatory cytokine responses induced by carbon-based nanomaterials in established *in vitro* models and *in vivo* lung. AMs GTR, BM-AMs, and MH-S cells were exposed for 24 h to MWCNT (16, 32, and 64 μg/ml), DWCNT (50, 100, and 200 μg/ml), or CNP (50, 100, and 200 μg/ml). Treatment with 0.1% Triton X-100 served as a positive control for cell lysis. Mice were exposed to CNP, DWCNT, or MWCNT as described above, and BAL fluid was collected 12 h after exposure for cytokine analysis. **(A)** Heatmaps showing cytokine and chemokine responses in the different *in vitro* AM models and BAL fluid. Values are presented as log2 fold changes relative to the corresponding untreated control. **(B)** Dot plot showing cytokine and chemokine gene expression in *in vivo* AMs derived from scRNA-seq analysis 12 h after CNP, DWCNT, or MWCNT exposure, for all in BAL (Fig. 4A) more than 2-fold increased proteins. Dot size represents the proportion of AMs expressing each gene, whereas color indicates relative expression level. Hierarchical clustering of the selected genes is shown on the right.

To assess how closely these *in vitro* responses reflected the pulmonary inflammatory response *in vivo*, the pattern were compared to that of mice intratracheally exposed to CNP (20 μg), DWCNT (50 μg), or MWCNT (15 μg), with bronchoalveolar lavage fluid collected after 12 h for cytokine analysis.^18^ Consistent with the *in vitro* findings, MWCNT induced the most pronounced cytokine response in bronchoalveolar lavage fluid (BAL), whereas DWCNT and CNP caused substantially weaker changes (**Fig. 4A**). We next used our scRNA-seq dataset to determine which of the 13 cytokines showing at least a twofold increase in BAL after CBN exposure, were also transcriptionally induced in AMs ((**Fig.4B**).^29^ Interestingly all 13 cytokines showed an induced pattern in AMs 12 h after particle exposure, but the moderate fold change levels and low proportion of the cytokine mRNA expressing cells need to be considered. However, overlapping lung AM gene expression with the cytokine release pattern from the three AM models, detect CCL2, CCL7 CXCL1 and IL-6 in both systems. *In vitro*, CCL7 is primarily released in BM-AM upon MWCNT exposure, but to a weaker extent also upon DWCNT and CNP, which matches the lung AM gene expression best for DWCNT. CCL2 was likewise released mainly by BM-AMs after MWCNT and DWCNT treatment, consistent with the lung data. In contrast, CXCL1 was more abundantly released by AMs GTR, mainly upon MWCNT or high doses of CNP. In lung AMs, however, elevated *Cxcl1* expression was limited to CNP exposure only.

Nevertheless, because BAL cytokines can originate from multiple pulmonary cell populations, these data provide complementary evidence but do not directly demonstrate that the measured mediators are derived from resident AMs. Additionally, not all cytokines need to be released following previously induced gene expression and some, in particular alarmins, reside in a preformed state in the cells to get rapidly released upon stimulation. To assess constitutively expressed cytokines and determine which were most reproduced by the three AM-models, cell lysates were analyzed using a 31-plex Bio-Plex cytokine assay. Interestingly only IL-16, a pro-inflammatory cytokine known to drive airway inflammation was abundantly detected in the primary AM models, and CXCL-16 and CCL-20 in contrast in MH-S lysates.

In general, both AMs GTR and BM-AMs reproduced the overall material-dependent inflammatory response pattern observed *in vivo* at best, with MWCNT consistently producing the strongest response. BM-AMs exhibited a broader inflammatory program and thus greater similarity to the BAL cytokine profile, suggesting that BM-AMs may better reproduce selected aspects of the acute inflammatory response of resident AMs to inhaled carbon-based nanomaterials.

### Carbon-based nanomaterial internalization and subsequent cellular organelle damage

A key function of resident alveolar macrophages is the continuous surveillance of the alveolar space and clearance of inhaled particles. To determine whether the investigated *in vitro* models reproduce these functional properties, we analyzed time-dependent cellular association of CBNs together with migratory activity in AMs GTR, BM-AMs, and MH-S cells. AM-models were exposed to low doses causing only moderate effects on cell viability (MWCNT: 8 μg/ml, DWCNT and CNP: 12.5 μg/ml).

Both primary AM models showed a rapid, pronounced time-dependent accumulation of MWCNT and CNP, with increased cell-associated particle signal peaking at around 6 h and slowly declining till 12 h, whereas BM-AM uptake was slower and increased till 12 h for MWCNT. DWCNT showed comparatively limited cell association (**Fig. 5A, B**). MH-S cells exhibited a distinct kinetic uptake profile, starting later and reaching for CNP highest levels 12 h after exposure. Consistent with their toxicity profile (**Fig. 3**), MWCNT reduced cell crawling of AMs GTR and BM-AMs, as assessed by measuring the two-dimensional migration trajectories, whereas migration was less affected in MH-S cells and even enhanced by DWCNT and CNP exposure (**Supplementary Fig. 4**).

**Fig. 5.**
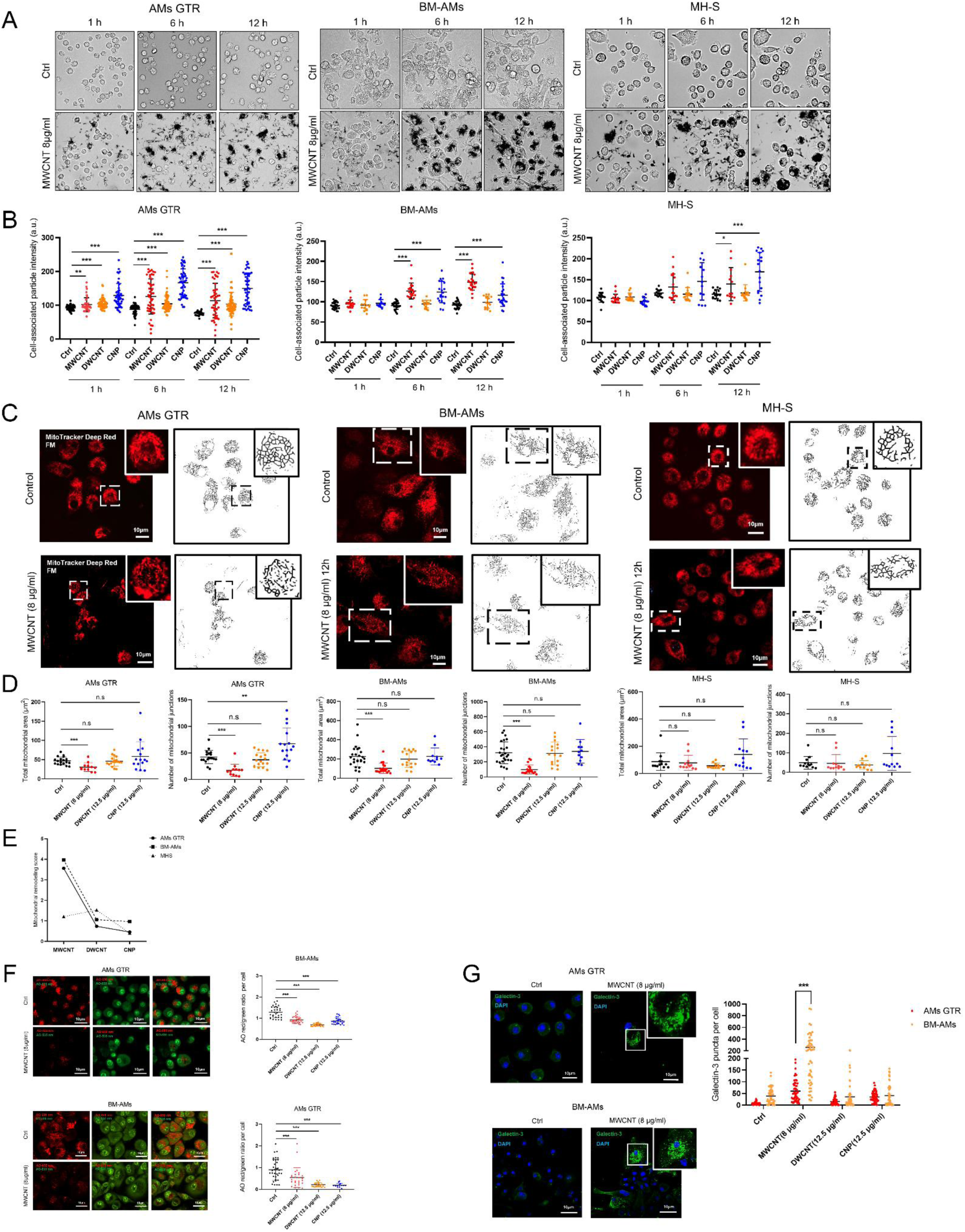
Carbon-based nanomaterial cell association and subsequent mitochondrial and lysosomal alterations in alveolar macrophage models. (A) Representative bright-field images of AMs GTR, BM-AMs, and MH-S cells following exposure to MWCNT (8 μg/ml) for 1, 6, and 12 h. (B) Quantification of cell-associated particle intensity in AMs GTR, BM-AMs, and MH-S cells following exposure to MWCNT (8 μg/ml), DWCNT (12.5 μg/ml), or CNP (12.5 μg/ml). Cell-associated particle signal was quantified from inverted bright-field images by measuring the mean pixel intensity within manually defined cellular regions of interest; higher values indicate greater particle-associated signal. a.u., arbitrary units. (C) Representative MitoTracker Deep Red images and corresponding skeletonized mitochondrial networks in untreated and MWCNT-exposed AMs GTR, BM-AMs, and MH-S cells after 12 h. Scale bars: 10 μm. (D) Quantification of total mitochondrial area and mitochondrial junction number following CBNs exposure. (E) Composite mitochondrial remodeling index derived from changes in mitochondrial area and junction number relative to untreated controls. (F) Representative acridine orange staining of AMs GTR and BM-AMs following CBNs exposure and quantification of the AO red/green fluorescence ratio per cell as an indicator of lysosomal acidification status. (G) Representative galectin-3 staining and quantification of galectin-3 puncta per cell in AMs GTR and BM-AMs following CBNs exposure. Galectin-3 is shown in green and nuclei in blue (DAPI). Scale bars: 10 μm. Data are presented as mean ± SEM from three independent experiments. Statistical significance was assessed by one-way ANOVA followed by Tukey’s multiple-comparisons test. \**P* < 0.05, \*\**P* < 0.01, \*\*\**P* < 0.001.

We next investigated whether these distinct particle-associated responses were accompanied by mitochondrial structural alterations. After 12 h exposure, MWCNT markedly reduced total mitochondrial area and mitochondrial junction number in both AMs GTR and BM-AMs, consistent with mitochondrial fragmentation and loss of network connectivity (**Fig. 5C, D**). Even though the MH-S model exhibited the lowest WST IC50 values for MWCNTs, no related mitochondrial shape changes could be detected for these cells. Summarizing the mitochondrial phenotype upon CBNs exposure by the Composite mitochondrial remodeling index (**Fig. 5E**) illustrates the great impact of MWCNTs particularly on the primary AM models. Next, lysosomal function was assessed by acridine orange (AO) staining. In AMs GTR and BM-AMs, exposure to all three CBNs decreased the red-to-green AO fluorescence ratio, indicating reduced lysosomal acidification (**Fig. 5F**). DWCNT exposure produced a mixed punctate staining pattern, suggesting partial alteration of lysosomal acidity. CNP aggregates noteworthy also caused pronounced local signal quenching, generating characteristic dark regions within the lysosomal compartment. Whereas MH-S cells exhibited a visually smaller lysosomal pool and distinct lysosomal morphology, including abnormal lysosomal fusion following DWCNT exposure (**Supplementary Fig. 5**).

To better distinguish lysosomal deacidification from lysosomal membrane permeabilization, galectin-3 puncta formation was additionally assessed to specifically detect damaged organelle membranes. Given the markedly smaller and morphologically distinct lysosomal compartment in MH-S cells, this analysis was restricted to AMs GTR and BM-AMs. MWCNT exposure induced a clear increase in galectin-3 puncta in both AMs GTR and BM-AMs, with a stronger response in BM-AMs (**Fig. 5G**). In contrast, DWCNT and CNP induced little or no detectable galectin-3 puncta formation under the same conditions, indicating that the pronounced lysosomal membrane damage observed in response to MWCNT was not reproduced by the other CBNs (**Supplementary Fig. 6**).^39^

Taken together, these findings demonstrate that the investigated AM models differ not only in particle- associated kinetics but also in the extent of mitochondrial and lysosomal injury. BM-AMs displayed the strongest coupling between particle exposure, organelle stress, and lysosomal membrane damage, whereas AMs GTR showed a similar but somewhat less pronounced response and MH-S cells exhibited only little organelle remodeling.

## Discussion

Alveolar macrophages (AMs) are essential for maintaining pulmonary homeostasis and responding to inhaled pathogens and pollutants, with their phenotype and function strongly shaped by the alveolar niche.^20,42–44^ Advances in single-cell transcriptomics have provided increasingly precise molecular benchmarks for defining AM identity and responses to inhaled toxicants.^45^ Although *in vitro* macrophage models are indispensable for mechanistic studies, controlled exposure experiments, and higher-throughput testing while reducing animal use, their ability to reproduce the phenotype and responses of AMs in vivo remains insufficiently defined.^46,47^ Here, we therefore systematically compared multiple *in vitro* AM models with *in vivo* AMs to determine their phenotypic and transcriptomic fidelity and their suitability for assessing carbon-based nanomaterial toxicity.

The first major distinction between the models is the balance between availability and fidelity to resident AM identity. Primary AMs most closely represent the *in vivo* situation but are severely constrained by their low yield, with approximately 0.2 million cells obtained per mouse. In contrast, AMs GTR and BM-AMs provide approximately 2 and 8 million cells per mouse, respectively, while MH-S and MPI offer essentially unlimited cell availability. Thus, increasing cell availability is accompanied by an increasing degree of deviation from the resident AM phenotype. However, freshly isolated AMs rapidly undergo functional adaptation once removed from the alveolar environment and placed in culture.^48^ Consequently, we recognized the loss of CD11c in BAL AMs 3 h after isolation *in vitro*, partially recovering after 24 h. The loss of CD11c was interestingly accompanied by an abnormal proinflammatory state. As evidenced by the fact that *ex vivo* cultured AM reacquired full transcriptional and epigenetic identity after transfer back into the lung.^48^ The application of the GTR cocktail, representing factors essential for AM differentiation and maintenance, is able to reconstruct the environmental *in vivo* cues. Our data indicated that this approach is sufficient for BAL AMs to inherit most of the transcriptome signature of *in vivo* AMs (> 65%) and its surface marker phenotype. In agreement, previous research also demonstrated GTR supplement supports the differentiation of bone marrow cells towards AMs, termed as BM-AMs.^37^ We recognized a coexistence of high levels of CD11b and CD11c with intermediate SiglecF, indicating that BM-AMs still preserve a monocyte-like footprint. In contrast, both MH-S and MPI share less than 50% of transcriptome similarity and exhibit low levels of CD11c and SiglecF. It should be noted that our 2D culture systems do not fully reproduce the metabolic cues of the alveolar niche, particularly continuous exposure to pulmonary surfactant, the relatively high alveolar oxygen tension at low glucose levels and respective cell-cell communications. The absence of these niche-specific signals may contribute to the pronounced alterations in oxidative phosphorylation and fatty acid metabolism observed across all *in vitro* models. An instructive role of pulmonary surfactant has been demonstrated previously, where Infasurf, a natural bovine surfactant containing lipids and the hydrophobic surfactant proteins SP-B/SP-C, together with GM-CSF, TGF-β, and IL-10, promoted the differentiation of human blood monocytes toward an AM-like phenotype characterized by enhanced lipid and cholesterol metabolic programs.^39^ However, commercially available animal derived surfactant can differ substantially in lipid concentration and surfactant protein composition and may exhibit batch variation, thus unable to fully reproduce the composition of naive alveolar surfactant.^49^ Moreover, the exogenous surfactant can actively modulate the inflammatory states of AMs, further limiting their usage in AM culture system.^50^ More physiological relevant approaches may involve co-culture of AMs with alveolar epithelial cells which secrete naive surfactant and GM-CSF. Recently proposed air liquid interface (ALI) AT2 and AM co-culture system demonstrated an elegant way to reproduce the alveolar niche especially epithelial cells-AM crosstalk.^51^ The unique alveolar niche, characterized by limited glucose availability, renders AM preferentially dependent on fatty acid oxidation rather than glycolysis.^52^ Accordingly, glucose availability should be considered when establishing *in vitro* AM culture systems. In the present study, we used low-glucose DMEM containing approximately 5.56 mM glucose. Although this concentration remains substantially higher than that reported for the alveolar lining fluid, the transcriptomic profiles did not indicate a dominant shift toward glycolysis in the long-term primary AM models. This suggests that differences in glucose availability alone are unlikely to account for the broader metabolic divergence observed between *in vitro* and resident AMs. Instead, the absence of additional niche-specific cues, including continuous exposure to pulmonary surfactant, the relatively high alveolar oxygen tension, and interactions with neighboring epithelial cells, may all contribute to the metabolic adaptations observed under *in vitro* conditions.

AM models are often used to test respiratory toxicity of various materials, and in this context the detrimental effects of carbon-based nanomaterials (CBNs), particularly fiber-shaped MWCNT on AMs have been widely reported.^53–56^ Consistent with these findings, our study demonstrated a similar dose-dependent cytotoxic response to MWCNT, DWCNT and CNP exposure in *in vivo* AMs, AMs GTR, and BM-AMs. Although MH-S cell viability clearly reproduced the *in vivo* toxicity ranking (MWCNT > DWCNT > CNP), it is unclear how much this correlation may be attributed to a particle mediated inhibition of proliferation, since little agreement with cytotoxicity (LDH release) was observed. The sequential events of AM mobilization, particle phagocytosis, and subsequent organelle damage represent some of the earliest cellular responses to CBNs exposure, preceding overt cell death and inflammatory activation. After inhalation and depositing on the alveolar surface, inhaled CBNs get sensed by a yet unknown mechanism and rapidly internalized by AMs.^18^ In this context, we revealed that AMs GTR and BM-AMs effectively internalized MWCNT and DWCNT as early as 6 h after exposure, whereas MH-S showed a somewhat delayed internalization of both CBNs. This delayed uptake may be related to the heterogeneous growth pattern of MH-S cells, with approximately 20% of the cells remaining in suspension and 80% growing adherently. For fibrous materials the interaction with macrophages is strongly impacted by shape (aspect ratio) and fiber stiffness. Recent work by Broßell et al. showed that NR8383 rat AMs can bend and internalize sufficiently flexible long fibers, whereas fibers with higher rigidity are likely to evoke frustrated phagocytosis.^36^ These findings are consistent with previous reports that long, rigid MWCNT induce frustrated phagocytosis and impair macrophage function, potentially through disruption of cytoskeletal dynamics and reduced cell motility.^57,58^ While upon exposure to surfactant dispersed MWCNT, only AM death but not the for frustrated phagocytosis prototypic IL-1b release was detected in our *in vivo* and *in vitro* studies,^29^ nevertheless, this would explain the decreased cell motility triggered by MWCNT in both AMs GTR and BM-AMs but for some reason not in semi-adherent MH-S cells. We also found that despite their micro-sized agglomerates, all three models were to some extent capable of internalizing DWCNT or exhibit membrane association. This is in line with previous *in vivo* data from mice, where DWCNT uptake by Mac3/LAMP2+ airspace macrophages was reported.^24^ Despite substantial CNP uptake in all three cell models, cell migration speed only increased in MH-S cells which is in contrast with previous *in vivo* findings showing that CNP actually boosted AM motility in the alveoli.^18^ The discrepancy between the intravital data and our in vitro migration assay suggests that CNP-induced AM motility is context dependent and that migration on a culture surface may involve mechanisms distinct from those operating on the alveolar epithelium *in vivo*, where LFA-1/ICAM-1 interactions have been implicated.^16^

Phagocytosed particles finally reach the lysosomal compartment, but besides representing the stomach of the cell, the lysosome serves as a signaling hub that integrates phagocytosis, autophagy, extracellular vesicle trafficking and mitochondrial activity, thereby influencing both cell fate - survival or death - and the fate of internalized particles - whether they are retained or exocytosed.^59–62^ Previous work from Kokot et al., have demonstrated that intracellular quarantining and intercellular cycling of poorly degradable metal- oxide nanomaterials can contribute to persistent pulmonary inflammation, underscoring the importance of intracellular particle fate.^6^ This mode of action, however, has not been observed for CBNs. The ability of MWCNT to disrupt lysosomal integrity, resulting in subsequent cathepsin B release in AMs *in vivo* is seen as a central feature of fiber toxicity.^40^ In our *in vitro* model, both AMs GTR and BM-AMs but not MH-S recapitulated this lysosomal impairment by MWCNT. Lysosomal rupture and membrane permeabilization have recently been shown by proteomic profiling of NR8383 AMs exposed to nanofibers, including Mitsui- 7 MWCNT.^54^ Notably, that study identified extracellular release of multiple lysosomal hydrolases, including several cathepsins, while membrane-associated lysosomal proteins remained largely unchanged, thereby providing additional evidence that lysosomal dysfunction represents an early mechanistic consequence of nanofiber–macrophage interactions.^54^ In our study, DWCNT and CNP exposure additionally reduced lysosomal acidity in AMs GTR and BM-AMs, potentially reflecting a disruption of the lysosomal acidification or related v-ATPase function. Lysosomal disruption and subsequent leakage of cathepsins can further impair mitochondrial function.^63^ Consistent with previous *in vivo* observations of decreased mitochondrial total mass upon MWCNT exposure,^41^ we confirmed that both mitochondrial mass and junctions are decreased in AMs GTR and BM-AMs (**Fig. 5C, D**). Of which, BM-AMs are more sensitive possibly due to a higher mitochondria number (**Fig. 5E**). The unaffected mitochondrial structure in both AMs GTR and BM-AMs following DWCNT and CNP exposure matches with the higher mass and surface doses required *in vivo* to cause equal levels of acute inflammation as compared to MWCNT.^20^ Although previous *in vivo* studies have reported mitochondrial ultrastructural changes in AMs following CNP exposure, these findings were largely obtained under chronic and extremely high-burden exposure conditions,^64^ eventually leading to overload and related to effects described here for excessive in vitro doses > 100 μg/ml (**Fig. 3B**).

Damaged organelles and particular mitochondria and lysosomes induce cellular stress, inflammation and cell death. In line with this, we have proposed MWCNT to trigger necroptotic AMs cell death as driver of chronic lung inflammation in mice.^20^ The identification of those pro-inflammatory cytokines released into the airspace and recovered by BAL, that originated from AMs, is technically challenging. To approach this, we combined BAL cytokine data from CBNs exposed mice with corresponding single cell RNA sequencing data from their lungs. From the identified 13 cytokines elevated in BAL by CBNs and associated with proinflammatory signaling, 10 were in AMs also induced at the gene expression level and in an exposure-specific manner (**Fig. 4B**). Out of these only the release of two cytokines, CCL2 and CCL7, could be reproduced at the *in vitro* level for DWCNT and MWCNT exposed AMs GTR and BM-AMs. No cytokine response could be reproduced in MH-S cells and CNP exposure failed to cause cytokine release in any of the models. While the latter agrees with previous studies showing no acute proinflammatory AM stimulation by CNPs,^20,22^ the lack of MH-S responsiveness questions the suitability of this model for investigations about the pro-inflammatory potency of CBNs.

A final question for this comparative study is of course an evaluation or ranking of the performance of the different tested AM models and we have summarized this in **Table. 2**. While the use of cell lines has clear advantages regarding the availability of high cell numbers, their immortalization and *in vitro* adaption seems to go at the expense of AM identity and responsiveness. Yet AMs GTR and BM-AMs, which still exceed the limited cell availability of primary AMs, show relatively high AM identity, but still differ in basic metabolic pathways. For that reason, the scientific and sound application of any *in vitro* model depends on the selection of the respective responses to be modeled. For the AM models tested here, BM-AMs seem to work best to study the inflammatory and cytotoxic response to CBNs and likely other poorly soluble particles of low toxicity. BM-AMs may similarly be well suited for assays related to fiber toxicity. In contrast, and because of their proliferative capacity allowing high throughput testing approaches of different nanomaterials, MH-S have proven to represent a well-suited model.^6^

## Conclusion

In summary we show a broad comparison of different murine AM models with mouse AMs at a descriptive and toxicological-functional level. While it is difficult to identify one model for all demands, our data indicate a clear compromise in terms of true-to-life physiology for the use of cell lines but at the same time also show that already the isolation of AMs and their culture at conventional submerged conditions, causes artificial reprogramming. Due to its proliferative capacity, the MH-S cell line might have a rightful place in cell viability high throughput applications, but the inflammatory responsiveness to CBNs seems inadequate. If the additional effort allows, the use of primary bone macrophages derived BM-AM is advisable, since this model performed best concerning basal AM phenotype, and cytotoxic and inflammatory response to the test particles.

## Materials and reagents

Spherical carbon nanoparticles (CNP; Printex90, Degussa, Frankfurt, Germany), double-walled carbon nanotubes (DWCNT; Nanocyl 2100, Auvelais, Belgium) and multi-walled carbon nanotubes (MWCNT; Mitsui-7, Hodogaya Chemicals, Mitsui, Japan) were used in this study as described previously (ACSnano 2025).

Prior to cell exposure, CBNs were dispersed in whole-lung pulmonary surfactant isolated from porcine lungs.^65^ Even if sterile conditions might be difficult to accomplish during the early steps of surfactant isolation, surfactant concentrations up to 200 μg/ml did not trigger any detectable inflammatory activation, such as IL-1a or IL-1b release from BAL AMs. NP and DWCNT stock suspensions (2 mg/ml) were diluted in low glucose DMEM (Thermo Fisher Scientific, Cat. No. 11885084; 1 g/L glucose) to a working concentration of 200 μg/ml and serially diluted two-fold to 100, 50, 25, and 12.5 μg/ml. MWCNT stock suspension (1.8 mg/ml) was diluted in complete culture medium to 64 μg/ml and serially diluted to 32, 16, 8, and 4 μg/ml.

### Cell line culture

The murine alveolar macrophage cell line MH-S was obtained from the American Type Culture Collection (ATCC; CRL-2019). MPI cells, originally generated from fetal livers of 15–19-day-old mouse embryos, were cultured as previously described.^34^

MH-S cells were maintained in RPMI 1640 medium supplemented with 10% fetal bovine serum (FBS), 100 U/mL penicillin, 100 μg/mL streptomycin, 0.1% 2-mercaptoethanol, and 1% glutamine. MPI cells were cultured in the same basal medium supplemented with murine GM-CSF at 30 ng/ml. Cells were maintained at 37 °C in a humidified incubator containing 5% CO_2_.

### Isolation and *in vitro* culture of primary alveolar macrophages

Mice were anesthetized by intraperitoneal administration of a ketamine/xylazine mixture containing ketamine (150 mg/kg body weight) and xylazine (10 mg/kg body weight). Procedures were initiated after adequate depth of anesthesia had been confirmed by the absence of reflex responses. Animals were subsequently euthanized by exsanguination via blood withdrawal from the inferior vena cava under deep anesthesia.

Bronchoalveolar lavage (BAL) was performed by repeated instillation and gentle aspiration of 1 ml ice-cold sterile Ca^2+^/Mg^2+^-free phosphate-buffered saline (PBS) through a tracheal cannula for 10 cycles, yielding approximately 10 ml BAL fluid per mouse. BAL fluid was centrifuged at 400 × g for 10 min at 4 °C, and the resulting cell pellet was resuspended in complete culture medium.

Cell numbers and viability were determined by trypan blue exclusion. For long-term differentiation, primary AMs were cultured for 7 days in complete medium supplemented with murine GM-CSF (20 ng/ml), human TGF-β (2 ng/ml), and the PPARγ agonist rosiglitazone (0.1 μM). The culture medium was replaced every 3 days.

For detachment, cells were washed once with prewarmed PBS and incubated with prewarmed Accutase for 30 min at room temperature. Complete medium was added to terminate the reaction, and cells were gently resuspended by pipetting. Cell suspensions were centrifuged at 400 × g for 5 min at 4 °C and resuspended in the appropriate buffer or culture medium for subsequent experiments.

### Bone marrow cell isolation and in vitro differentiation

Female C57BL/6J mice (8-12 weeks old) were purchased from Charles River (Sulzfeld, Germany) and housed in individually ventilated cages with food and water. All animal experiments were approved by the local Animal Care and Use Committee (District Government of Upper Bavaria, approval number ROB- 55.2-2532.Vet_02-15-67). Mice were euthanized in accordance with previously described protocols. Hind limbs were removed and transferred to a sterile dish containing ice-cold buffer solution (PBS without Ca2+/Mg2+ supplemented with 2% FBS and 2 mM EDTA). Bone marrow was harvested by flushing each bone with a 5–10 ml cold buffer using a 10 ml syringe into a 50 ml conical tube kept on ice. The suspension was passed through a 70 μm cell strainer (Miltenyi Biotech) into a new chilled tube to remove bone chips and debris, then centrifuged at 400 × g for 5 min at 4 °C. Red blood cells were lysed by red blood lysis buffer (Thermo Fisher Scientific). The cell pellet was collected and counted with 0.2% Trypan blue dye (Sigma Aldrich,Taufkirchen, Germany) and seeded to 20 ml culture flasks supplemented with GM-CSF (20 ng/ml) and TGF-β (2 ng/ml) for the first 7 days. The medium was then replaced with GM-CSF (20 ng/ml), TGF-β (2ng/ml), and the PPARγ agonist rosiglitazone (0.1 μM) for an additional 2 days. At day 9, cells were detached following the same protocol described above. Cell pellets were resuspended and cell numbers were calculated again. For live cell imaging, 1 × 10^5^ cells were seeded into 24 well black plates (ibidi, Cat.No:82426).

### Flow cytometry

Cells were detached and resuspended in a density of 10×10^6^ cells/ml in flow cytometry buffer. Cells were incubated with SiglecF-APC (anti-mouse, Cat. No. 130-112-333, Miltenyi Biotec), CD11b-PE (anti-mouse, Cat. No. 130-113-235, Miltenyi Biotec), and CD11c-PE Cy5.5 (anti-mouse, Cat. No. 45-0114-80) antibodies (1:50) for 15 min on ice. The flow cytometric analysis was performed on a BD FACSCanto II flow cytometer (BD 830 Biosciences) with BD FACSDiva v6.1.3 software.

### Bulk RNA sequencing

Total RNA was isolated from AMs 3 h and 24 h, AMs GTR, BM-AMs, MH-S, and MPI cells using the RNeasy Plus Mini Kit (Qiagen) according to the manufacturer’s instructions. RNA concentration and purity were assessed using a NanoDrop spectrophotometer (Thermo Fisher Scientific).

RNA-seq libraries were prepared using the Illumina Stranded mRNA Prep, Ligation kit (Illumina, cat. no. 20040534) according to the manufacturer’s protocol. Briefly, poly(A)-containing mRNA was isolated using oligo(dT) magnetic beads, fragmented, and reverse-transcribed into strand-specific cDNA. Following second-strand synthesis, cDNA fragments were adenylated, ligated to indexed adapters, purified using AMPure XP beads, and PCR-amplified to generate dual-indexed libraries.

Final libraries were assessed using an Agilent 2100 Bioanalyzer and subsequently pooled for paired-end sequencing on an Illumina sequencing platform. Raw sequencing reads were processed to obtain gene- level read counts, which were used for downstream transcriptomic analyses.

### Data analysis and visualization

Transcriptomic analyses were performed using Python 3.11 with pandas, NumPy, SciPy, scikit-learn, Matplotlib, and Seaborn, and R for differential expression and pathway analyses. For comparison of bulk RNA-seq and scRNA-seq datasets, low-expressed genes were filtered out and expression values were normalized using the housekeeping genes *Hprt* and *Gapdh*. PCA, Pearson correlation analysis, and heatmaps were generated to assess transcriptomic similarity between in vitro AM models and resident AMs. Differentially expressed genes were identified using the limma-voom workflow with edgeR and limma, using adjusted P < 0.05 and |log2FC| > 1 as significance thresholds. GSEA was performed in R v4.5.0 using clusterProfiler v4.16.0, and enrichment was expressed as normalized enrichment score (NES).

### WST-1 assay

Cells were seeded in 24-well plates at 1 × 10^5^ cells/well in 500 μl complete culture medium and incubated for 24 h at 37 °C. Cells were subsequently exposed for 24 h to CNP or DWCNT at 12.5, 25, 50, 100, or 200 μg/ml, or to MWCNT at 4, 8, 16, 32, or 64 μg/ml.

Following exposure, culture medium was replaced with 300 μl fresh culture medium containing 30 μl WST- 1 reagent (Cell Proliferation Reagent WST-1; Roche Diagnostics, Mannheim, Germany; cat. no. 11644 807001). Cells were incubated for 30 min, after which the supernatant was collected and centrifuged at 14,000 rpm for 10 min to remove particulate material. Absorbance was measured at 440 nm using a Tecan microplate reader.

Relative cell viability was calculated as: Cell viability (%) = [(OD_sample − OD_blank) / (OD_control − OD_blank)] × 100.

### LDH assay

Cells were seeded at 1 × 10^5^ cells/well in 24-well plates containing 500 μl culture medium and incubated for 24 h before CBNs exposure. Cells were treated with the same concentration ranges of CNP, DWCNT, and MWCNT used for the WST-1 assay.

A Triton X-100-treated group served as the high-release control. Following exposure, supernatants were collected and centrifuged at 14,000 rpm for 10 min to remove residual particles. Aliquots of supernatant were transferred to a 96-well plate and incubated with LDH reaction mixture (Roche Diagnostics; cat. no. 11644793001) according to the manufacturer’s instructions. Absorbance was measured at 492 nm using a Tecan microplate reader.

Relative LDH release was calculated as: Cytotoxicity (%) = [(OD_sample − OD_blank) / (OD_high control – OD_blank)] × 10

### Immunofluorescence staining

Cells were seeded at 5 × 10^4^ cells/well onto #1.5 glass coverslips in 24-well plates and incubated for 24 h. Following exposure to the indicated CBNs, cells were washed three times with PBS and fixed with 4% paraformaldehyde for 20 min at room temperature.

Cells were permeabilized and blocked for 1 h in PBS containing 5% goat serum and 0.3% Triton X-100. Samples were subsequently incubated overnight at 4 °C with anti-Galectin-3 primary antibody (clone A3A12; Invitrogen/Thermo Fisher Scientific, cat. no. MA1-940) diluted 1:50 in antibody diluent containing 1% BSA and 0.3% Triton X-100.

Following washing, cells were incubated for 1 h with the appropriate fluorescent secondary antibody (Goat anti-Rabbit IgG (H+L) Alexa Fluor 488, A11008, Thermo Fisher Scientific, Waltham, MA USA) and counterstained with DAPI. Samples were mounted using DAKO fluorescence mounting medium (Dako Omnis, Agilent, Santa Clara, USA) and imaged using a Zeiss LSM 880 confocal laser-scanning microscope driven by Zen 2.3 (“blue version”) software (Carl Zeiss).

Galectin-3 puncta were quantified using Fiji/ImageJ. Individual cells were manually outlined as regions of interest (ROIs), and the Galectin-3 fluorescence channel was background-corrected and thresholded using consistent settings across all images. Galectin-3-positive puncta within each cellular ROI were quantified using the Analyze Particles function and expressed as the number of puncta per cell. A total of 48-53 individual cells per condition, pooled from three independent experiments, were analyzed.

### Mitochondrial staining

Cells were seeded in a black 24-well plate with glass bottom at a density of 100,000 cells per well and incubated at a 37 °C humidified incubator (95% humidity, 5% CO_2_) for 24 h. Cells were stained with MitoTracker Deep Red FM (Thermo Fisher Scientific; cat. no. M22426) at 200 nM for 15 min and washed with prewarmed culture medium before CBNs exposure. Cells were subsequently exposed for 12 h to MWCNT (8 μg/ml), DWCNT (12.5 μg/ml), or CNP (12.5 μg/ml). Live-cell images were acquired using a Zeiss Celldiscoverer 7 microscope with excitation/emission of 644 / 665 nm appropriate for MitoTracker Deep Red FM. Cells were exposed to MWCNT (8 μg/ml), DWCNT (12.5 μg/ml) and CNP (12.5 μg/ml) for 12 h and immediately imaged by using live cell confocal microscope (Zeiss celldiscoverer 7), equipped with 20× / 0.7 objective (Plan-Apochromat) and TL LED light source for laser-illumination (Pinhole: 1.00 AU / 75 μm, 640 nm:1.00%), Unidirectional Scanning microscope. The cell culture condition during live cell imaging was 5% CO_2_ and 95% humidity. Mitochondrial morphology was quantified according to the previously described pipeline.^66^

### Acridine Orange (AO) staining

Cells were seeded in a black 24-well plate (ibidi, Cat.No:82426) with glass bottom at a density of 100,000 cells per well and cultured in a 37 °C humidified incubator (95% humidity, 5% CO_2_) for 24 h. Cells were exposed to CNP (12.5 μg/ml), DWCNT (12.5 μg/ml) and MWCNT (8 μg/ml) for 24 h. Each well was exchanged with a complete culture medium containing 1 μg/ml of AO (A9231, SigmaAldrich, Germany) and incubated for 15 min. Images of lysosomes were acquired by using a live-cell confocal microscope (Zeiss cell discoverer 7) equipped with 50×/1.2 objective (Plan-Apochromat), bidirectional scanning and TL LED light source for laser-illumination for two channels: green (Ex/Em: 493 nm / 517 nm, 0.3% laser power, 1.00 AU / 64 μm pinhole) and red (Ex/Em: 553 nm / 568 nm, 0.1% laser power, 0.79 / 64 μm pinhole). The cell culture condition during live cell imaging was 5% CO_2_ and 95% humidity. Cells were outlined as ROI from the green channel, the quantification of AO in each cell was performed by subtracting background signal and calculating the ratio of red signal to green signal intensity by using imageJ/Fiji.

### Cell migration analysis

Cells were seeded in a black, glass-bottom 24-well plate (ibidi, Cat.No:82426) at a density of 10,000 cells per well and cultured at 37 °C in a humidified incubator (95% humidity, 5% CO_2_) for 24 h. Cells were treated with CNP (12.5 μg/ml), DWCNT (12.5 μg/ml) and MWCNT (8 μg/ml). The cell movement was tracked by staining cellular nuclei with hoechst 33342 (cat.62249, Invitrogen/Thermo Fisher Scientific) diluted at 1:1000 in culture medium. Observations were facilitated using a live-cell confocal microscope Zeiss Celldiscoverer 7 (CD7) equipped with 20×/0.7 objective (Plan-Apochromat), and TL LED light source for laser-illumination (Pinhole: 1.00AU/44 μm, 405nm:0.5%) for 12 h. The cell culture condition during live cell imaging was 5% CO_2_ and 95% humidity. Time-lapse images were acquired every 5 min and exported as AVI files.

Cell migration was analyzed using CellTraxx via the Fiji macro interface, as previously described by Børge Holme et al.^67^ Briefly, CellTraxx performs frame-to-frame X-Y drift correction, cell segmentation based on grayscale intensity, and centroid-based tracking of individual cells between consecutive frames using nearest-neighbor matching. Short interruptions in individual tracks were corrected by gap interpolation where applicable. Cell trajectories were used to calculate migration parameters, including velocity, accumulated distance, Euclidean distance, directness, and forward migration index. Crawling velocity was used as the primary migration parameter for statistical comparison between treatment groups.

For AMs GTR, 35 control cells, 36 MWCNT-exposed cells, 44 DWCNT-exposed cells, and 25 CNP- exposed cells were successfully tracked and included in the analysis. For BM-AMs, 414 control cells, 570 MWCNT-exposed cells, 659 DWCNT-exposed cells, and 486 CNP-exposed cells were tracked and included in the analysis. For MH-S, 91 control cells, 122 MWCNT-exposed cells, 118 DWCNT-exposed cells, and 100 CNP-exposed cells were tracked and included in the analysis.

### CBNs internalization quantification

Cells were seeded in a black glass-bottom 24-well plate (ibidi, Cat.No:82426) at a density of 10,000 cells per well and cultured at 37 °C humidified incubator (95% humidity, 5% CO_2_) for 24 h. Cells were subsequently exposed to MWCNT (8 μg/ml), DWCNT (12.5 μg/ml), or CNP (12.5 μg/ml) in a final volume of 500 μl per well. Cell-associated CBNs accumulation was monitored for 12 h using a Zeiss Celldiscoverer 7 in bright-field mode (Plan-Apochromat 20× / 0.95, TL LED light source, 20% light source intensity, unidirectional scan). The cell culture condition during live cell imaging is 5% CO_2_ and 95% humidity.

Images acquired at 1, 6, and 12 h were selected for quantitative analysis using Fiji/ImageJ. Individual cells were manually outlined as regions of interest (ROIs), and the images were inverted so that dark CBNs- associated regions appeared bright. The mean pixel intensity within each cellular ROI was calculated as the sum of pixel intensity values divided by the number of pixels within the ROI. Higher intensity values therefore indicated greater cell-associated CBNs signal. Values were expressed in arbitrary units (a.u.). Depending on the model, treatment, and time point, 14-73 individual cells were analyzed per condition. Specifically, 32-73 cells were analyzed for AMs GTR, 15-26 cells for BM-AMs, and 14-20 cells for MH-S.

### 31-plex assay

AMs GTR, BM-AMs and MH-S were treated with CNP (50, 100, 200 μg/ml), DWCNT (50, 100, 200 μg/ml) and MWCNT (16, 32, 64 μg/ml) for 24 h. The supernatant was collected and centrifuged for 10 min at 14,000 rpm. Cytokines and chemokines in each sample were quantified following the manufacturer’s protocol for the Bio-Plex Pro Mouse Chemokine Panel 31-Plex (Bio-Rad Laboratories, California, USA). Data acquisition was performed using a LuminexTM 200 plate reader (InvitrogenTM, Thermo Fisher Scientific, Carlsbad, USA), and the BioPlex Manager 6.2 software (Bio-Rad Laboratories, California, USA) was used for analysis. Standard curves were fitted using the logistic-5PL regression type. Heatmaps with hierarchical clustering were generated from z-score-scaled chemokine concentrations in R (v4.2.3) using pheatmap (v1.0.13) (https://cran.rproject.org/web/packages/pheatmap).

### Data availability

scRNA-seq raw data have been submitted to GEO (GSE273313), the bulk RNA-seq raw data is under submission. All other data supporting the findings of this study are available within the Article and Supporting Information.

### Statistical analysis

All data were shown as mean ± SEM and plotted by using GraphPad Prism 8 (GraphPad Software Inc., La Jolla, USA). Comparison between two or multiple groups was performed by using a two-sided Student’s t-test or One-way/Two-way ANOVA test. A statistical significance was defined as P value <0.05.

## Acknowledgments

The authors would like to particularly thank A. Fuchs and D. Kutschke for their excellent technical work supporting the project. The authors are grateful to the members of the animal facility of Helmholtz Munich for their professional support and assistance; Barbara Mosetter (Immunanalytik-Tissue Control of Immunocytes, IMA-TCI, Deutsches Forschungszentrum für Gesundheit und Umwelt, Helmholtz Zentrum München, Munich, Germany) for technical assistance with the Bioplex assay; A. Feuchtinger (Research Unit Analytical Pathology, Helmholtz Zentrum München, Munich, Germany) for support in live-cell and confocal imaging. We also acknowledge the technical support of Core Facility Genomics at Helmholtz Munich.

This work was supported by funding from the German Center for Lung Research (DZL), European Union (EU) Horizon 2020 Research and Innovation program (953183, HARMLESS) and European Union under the HORIZON-CL4–2022-DIGITAL-EMERGING-01 Program (GA101092741, NanoPass). H. Ren received a fellowship from China Scholarship Council (CSC, 202006240092). C. Voss acknowledges funding by the Deutsche Forschungsgemeinschaft (DFG, German Research Foundation)-Project 530370007.

## Author Contributions

H. Ren performed all the experiments and analyzed the data. L. Han and T. Stoeger contributed equally to this article as corresponding authors and supervised the work. H. Ren, L. Han and T. Stoeger designed and planned the entire study. M. Rehberg supervised cell trajectory analysis and all image quantifications. H. Ren drafted the manuscript. L. Han, M. Rehberg, C. Voss and T. Stoeger revised the manuscript.

