## Supplemental file for "Establishing toxicologically relevant primary alveolar macrophage models to enable a realistic assessment of nanomaterials safety"

1. Institute of Lung Health and Immunity (LHI), Comprehensive Pneumology Center (CPC), Helmholtz Munich, Neuherberg 85764, Germany; Member of the German Center for Lung Research (DZL), Munich 81377, Germany
2. Genomics Core Facility, Helmholtz Munich, Neuherberg 85764, Germany
3. Dept. Biochemistry and Molecular Biology, Faculty of Biology, and Research Institute "Hospital 12 de Octubre (imas12)", Complutense University, 28040 Madrid, Spain
4. Hannover Medical School, Clinic for Cardiac, Thoracic, Transplantation and Vascular Surgery, Leibniz Research Laboratories for Biotechnology and Artificial Organs (LEBAO), Hannover 30625, Germany; Biomedical Research in Endstage and Obstructive Lung Disease Hannover (BREATH), German Center for Lung Research (DZL), Hannover 30625, Germany
5. Suzhou Institute of Nano-Tech and Nano-Bionics (SINANO), Chinese Academy of Sciences, Suzhou 215123, China

**Contact information**

* Correspondence to T. Stoeger, L. Han

### Supplementary figures & legends

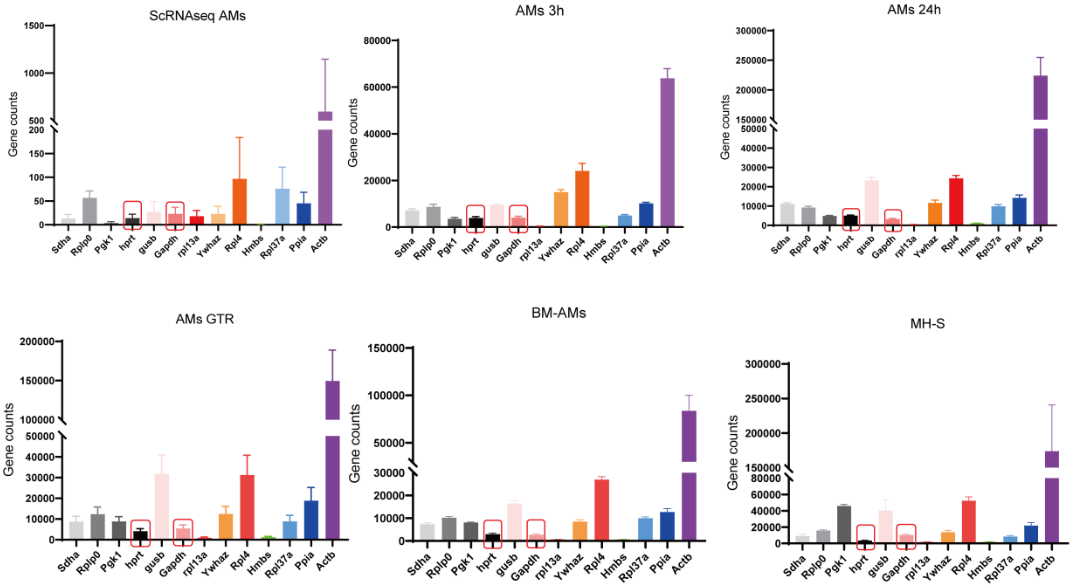


**Supplementary Fig. 1. Expression of candidate housekeeping genes across in vitro AM models**

DESeq2-normalized expression counts of 13 candidate housekeeping genes were evaluated across AMs 3 h, AMs 24 h, AMs GTR, BM-AMs, MH-S, and MPI cells. Bars represent the mean ± SEM of three biological replicates. *Hprt* and *Gapdh* (red boxes) were consistently detected across all models and, together with their reproducible expression across the scRNA-seq dataset, were selected as reference genes for cross-dataset normalization.

**Supplementary Table S1: List of 483 core alveolar macrophage genes used for PCA, correlation, and heatmap analyses (Excel file)**

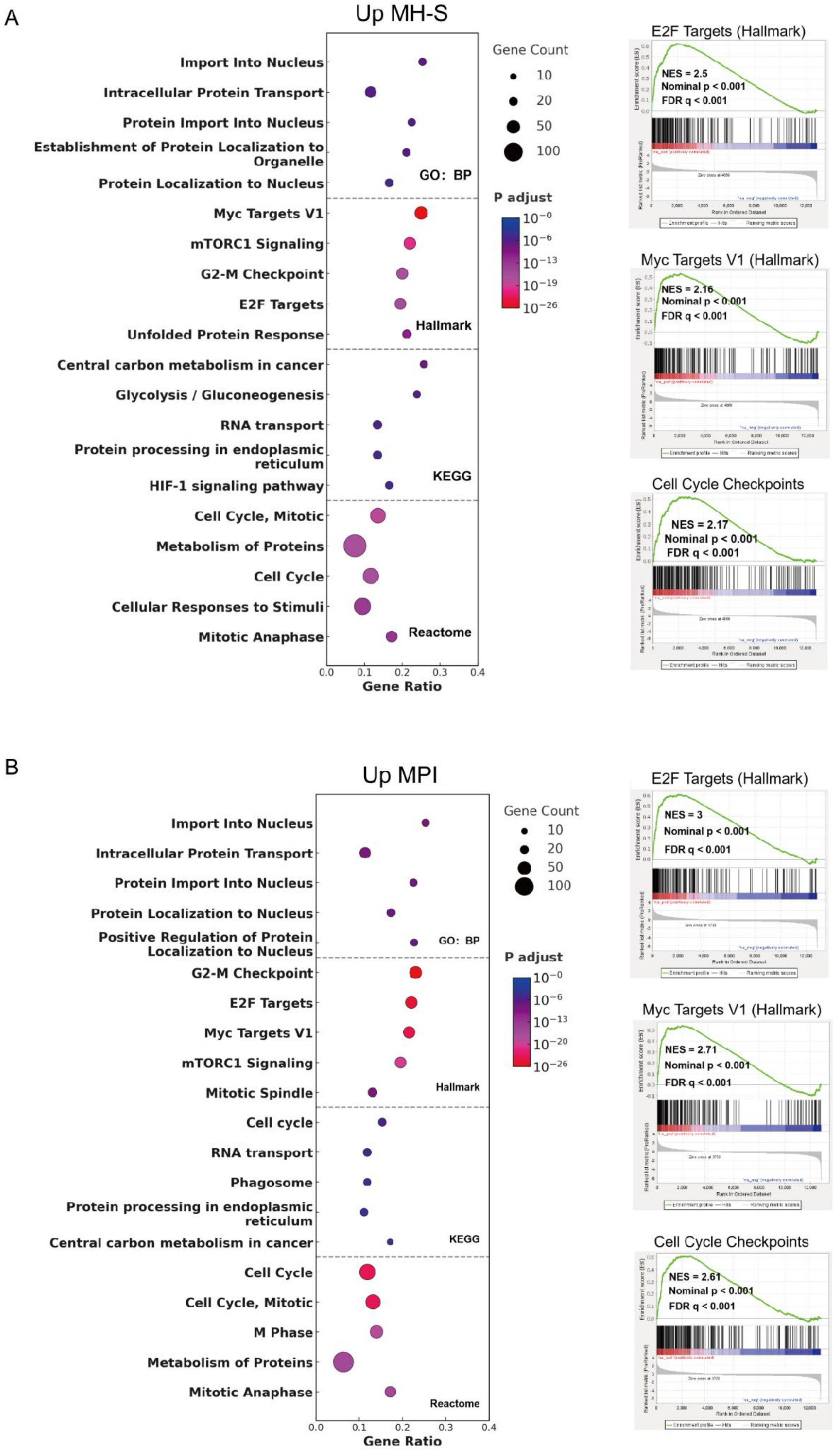
A
Up MH-S
Import Into Nucleus
Intracellular Protein Transport
Protein Import Into Nucleus
Establishment of Protein Localization to Organelle
Protein Localization to Nucleus
GO: BP
Myc Targets V1
mTORC1 Signaling
G2-M Checkpoint
E2F Targets
Unfolded Protein Response
Hallmark
Central carbon metabolism in cancer
Glycolysis / Gluconeogenesis
RNA transport
Protein processing in endoplasmic reticulum
HIF-1 signaling pathway
KEGG
Cell Cycle, Mitotic
Metabolism of Proteins
Cell Cycle
Cellular Responses to Stimuli
Mitotic Anaphase
Reactome
0.0 0.1 0.2 0.3 0.4
Gene Ratio
Gene Count
10
20
50
100
P adjust
$10^{-0}$
$10^{-6}$
$10^{-13}$
$10^{-19}$
$10^{-26}$
E2F Targets (Hallmark)
NES = 2.5
Nominal p < 0.001
FDR q < 0.001
Myc Targets V1 (Hallmark)
NES = 2.16
Nominal p < 0.001
FDR q < 0.001
Cell Cycle Checkpoints
NES = 2.17
Nominal p < 0.001
FDR q < 0.001
B
Up MPI
Import Into Nucleus
Intracellular Protein Transport
Protein Import Into Nucleus
Protein Localization to Nucleus
Positive Regulation of Protein Localization to Nucleus
GO: BP
G2-M Checkpoint
E2F Targets
Myc Targets V1
mTORC1 Signaling
Mitotic Spindle
Hallmark
Cell cycle
RNA transport
Phagosome
Protein processing in endoplasmic reticulum
Central carbon metabolism in cancer
KEGG
Cell Cycle
Cell Cycle, Mitotic
M Phase
Metabolism of Proteins
Mitotic Anaphase
Reactome
0.0 0.1 0.2 0.3 0.4
Gene Ratio
Gene Count
10
20
50
100
P adjust
$10^{-0}$
$10^{-6}$
$10^{-13}$
$10^{-20}$
$10^{-26}$
E2F Targets (Hallmark)
NES = 3
Nominal p < 0.001
FDR q < 0.001
Myc Targets V1 (Hallmark)
NES = 2.71
Nominal p < 0.001
FDR q < 0.001
Cell Cycle Checkpoints
NES = 2.61
Nominal p < 0.001
FDR q < 0.001

**Supplementary Fig. 2. Enrichment analysis of pathways upregulated in MH-S and MPI cells relative to resident alveolar macrophages**

**(A**) Functional enrichment analysis of genes upregulated in MH-S cells. **(B)** Functional enrichment analysis of genes upregulated in MPI cells. Bubble plots show the top enriched terms from Gene Ontology Biological Process (GO: BP), Hallmark, KEGG, and Reactome gene sets. The x-axis indicates gene ratio, bubble size represents the number of genes contributing to each enriched term, and bubble color denotes the adjusted P value. Representative gene set enrichment analysis (GSEA) plots for E2F Targets, Myc Targets V1, and Cell Cycle Checkpoints are shown on the right. Normalized enrichment score (NES), nominal P value, and false discovery rate (FDR) q value are indicated within each GSEA plot.

**Table S2. Physiochemical Characteristics and Dispersion Quality of CBNs.**

| Particle type | Hydrodynamic size (nm) | PdI | Dispersion in surfactant | Sedimentation in surfactant | Carbon (%) |
|---|---|---|---|---|---|
| MWCNT | 1314 | 0.364 | heterogeneous agglomerates | Clear time-dependent sedimentation | 98.1 |
| DWCNT | 951.6 | 0.824 | Highly polydisperse | Moderate-to-strong sedimentation | >90 |
| CNP(Printex90) | 2626 | 1.000 | Strong agglomeration | Strongest sedimentation | 99 |

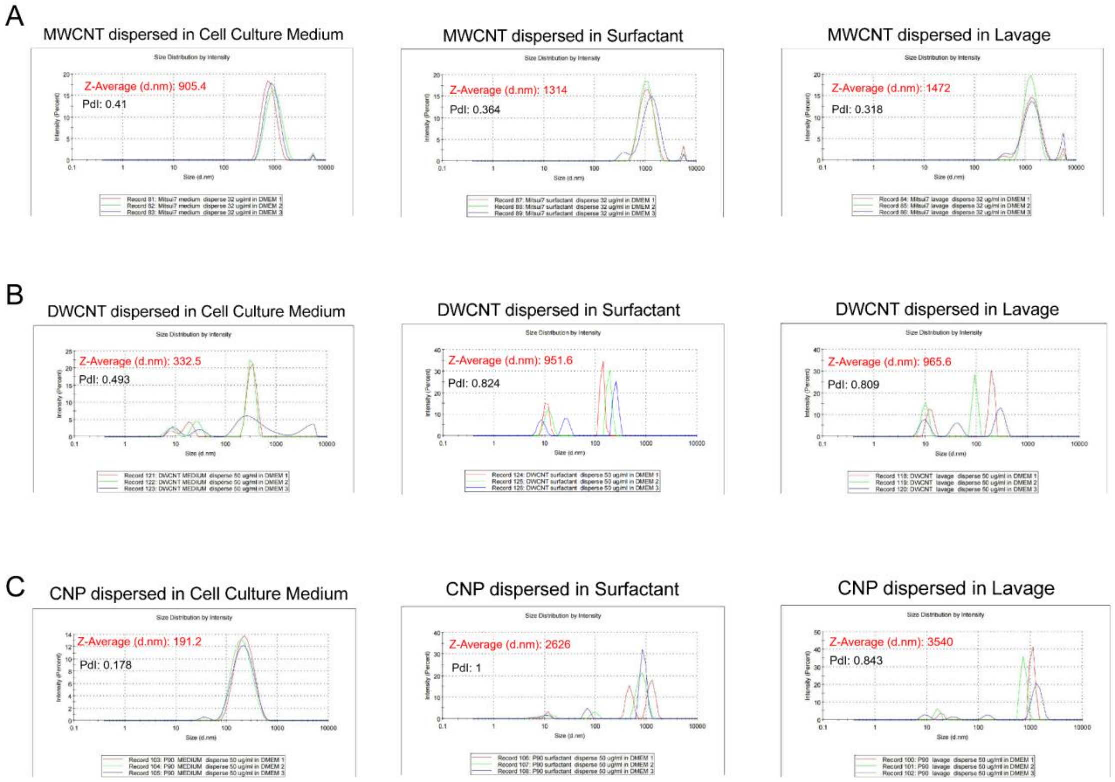


**Supplementary Fig. S3. Hydrodynamic size distribution of CBNs in different dispersion media**

Dynamic light scattering (DLS) analysis of **(A)** MWCNT, **(B)** DWCNT, and **(C)** CNP dispersed in cell culture medium, porcine surfactant, or bronchoalveolar lavage fluid. Size distributions are shown as intensity-weighted hydrodynamic diameter profiles from three measurements. The corresponding Z-average hydrodynamic diameter and polydispersity index (PDI) are indicated in each panel. MWCNT showed Z-average diameters of 905.4 nm, 1314 nm, and 1472 nm in cell culture medium, surfactant, and lavage fluid, respectively; DWCNT showed Z-average diameters of 332.5 nm, 951.6 nm, and 965.6 nm; and CNP showed Z-average diameters of 191.2 nm, 2626 nm, and 3540 nm.

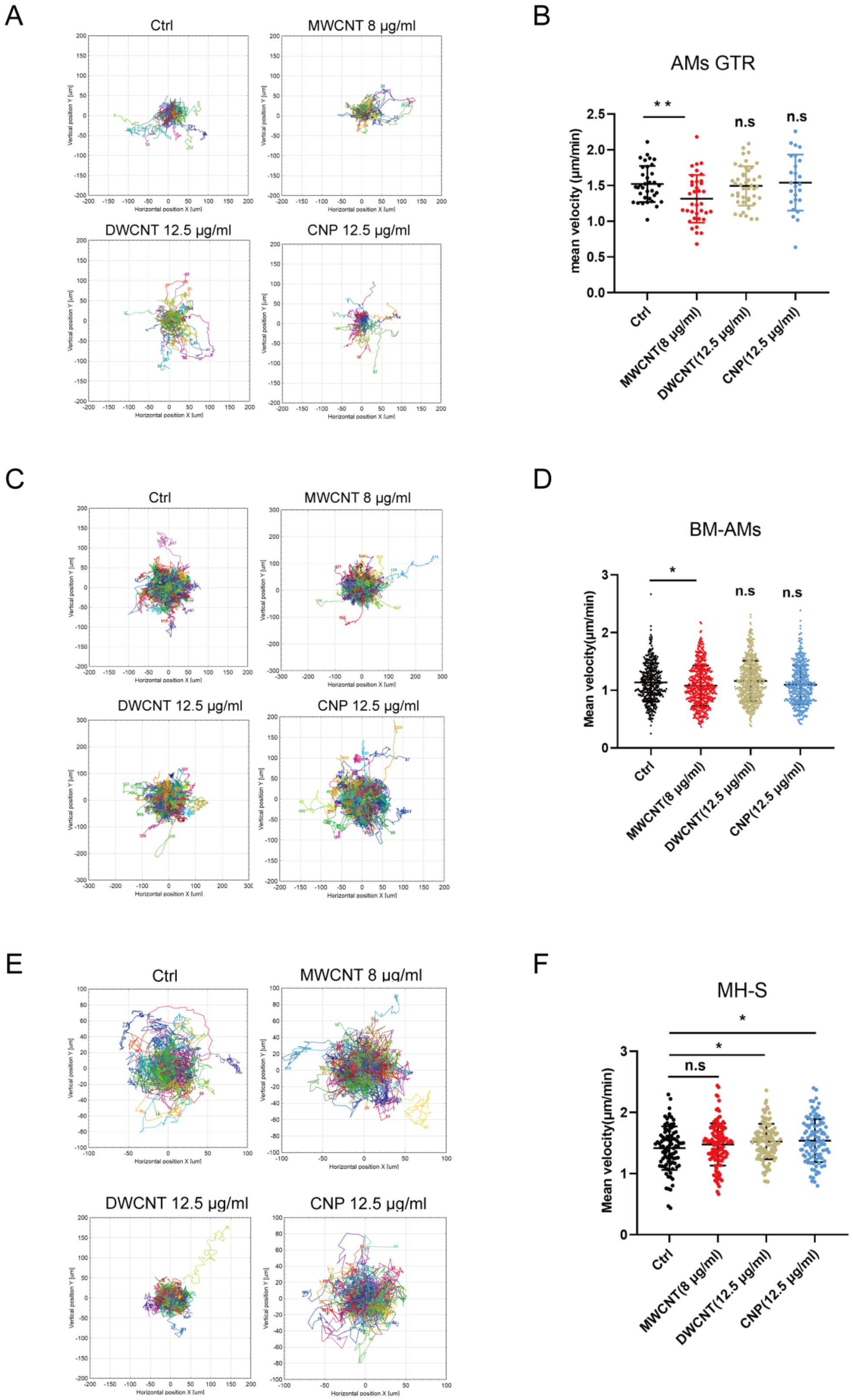
A
Ctrl
MWCNT 8 µg/ml
DWCNT 12.5 µg/ml
CNP 12.5 µg/ml
Horizontal position X [um]
Vertical position Y [um]
B
AMs GTR
mean velocity (µm/min)
**
n.s
n.s
Ctrl
MWCNT(8 µg/ml)
DWCNT(12.5 µg/ml)
CNP(12.5 µg/ml)
C
Ctrl
MWCNT 8 µg/ml
DWCNT 12.5 µg/ml
CNP 12.5 µg/ml
D
BM-AMs
Mean velocity(µm/min)
*
n.s
n.s
Ctrl
MWCNT(8 µg/ml)
DWCNT(12.5 µg/ml)
CNP(12.5 µg/ml)
E
Ctrl
MWCNT 8 µg/ml
DWCNT 12.5 µg/ml
CNP 12.5 µg/ml
F
MH-S
Mean velocity(µm/min)
*
*
n.s
Ctrl
MWCNT(8 µg/ml)
DWCNT(12.5 µg/ml)
CNP(12.5 µg/ml)

**Supplementary Fig. S4. Effects of CBNs exposure on macrophage migration**

**(A, C, E)** Representative two-dimensional migration trajectories of AMs GTR, BM-AMs, and MH-S cells, respectively, recorded over 12 h following exposure to control conditions, MWCNT (8 µg/ml), DWCNT (12.5 µg/ml), or CNP (12.5 µg/ml). Each colored line represents the migration path of an individual cell. **(B, D, F)** Quantification of mean migration velocity for AMs GTR, BM-AMs, and MH-S cells, respectively. Each dot represents one tracked cell, and horizontal lines indicate mean ± SEM. Statistical significance was assessed relative to the control group; $^*P < 0.05$, $^{**}P < 0.01$; n.s., not significant.

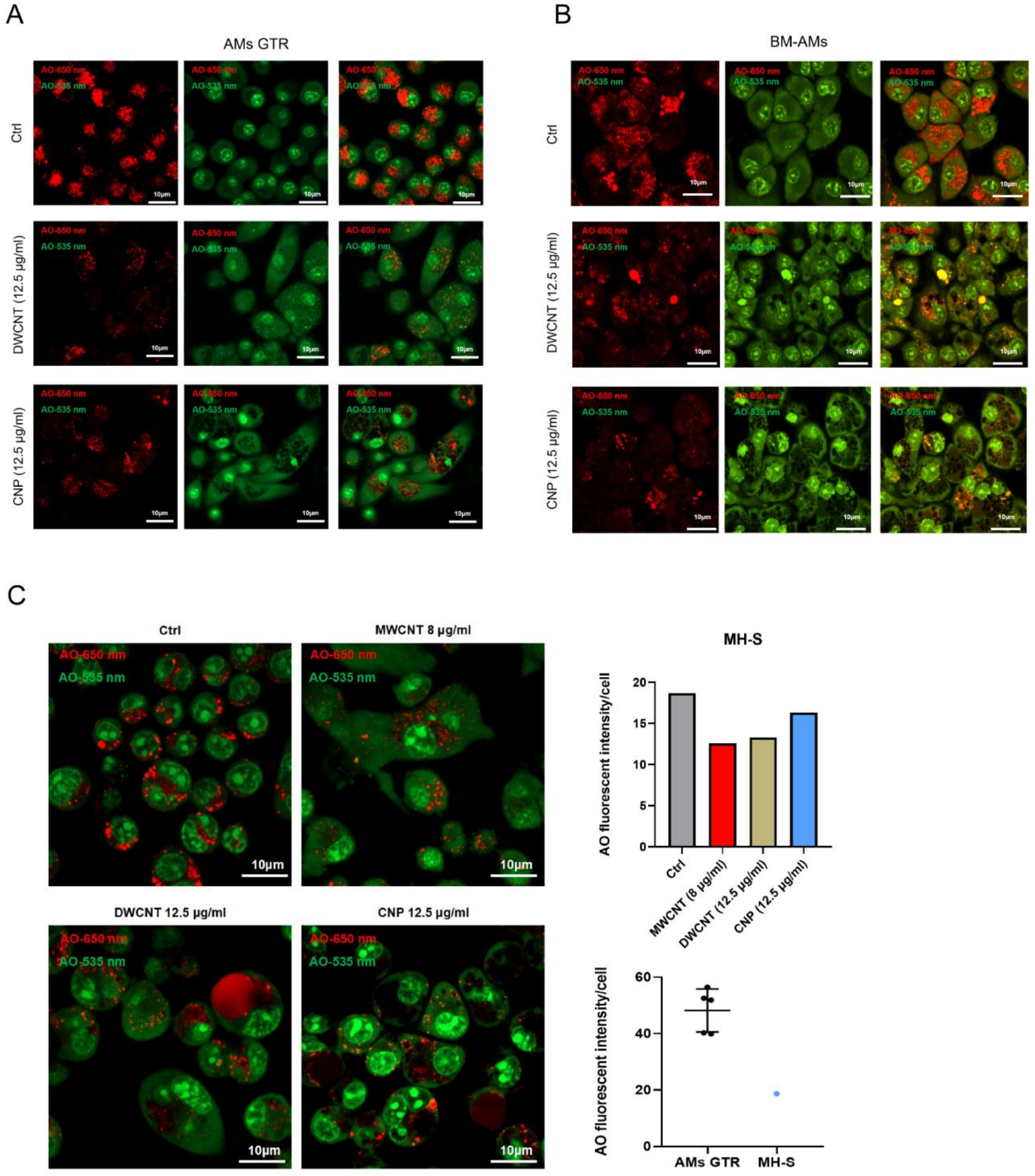


**Supplementary Fig. S5. Acridine orange staining reveals lysosomal alterations following CBNs exposure**

**(A, B)** Acridine orange staining reveals alterations in lysosomal acidification following exposure to DWCNT (12.5 μg/ml) or CNP (12.5 μg/ml) in AMs GTR and BM-AMs. The green monomeric form of acridine orange was detected at 535 nm, whereas the red dimeric form was detected at 650 nm; merged images are shown in the right column. Scale bars, 10 μm. (**C**) Representative AO fluorescence images of MH-S cells under control conditions or following exposure to MWCNT (8 μg/ml), DWCNT (12.5 μg/ml), or CNP (12.5 μg/ml), together with quantification of AO fluorescence intensity per cell. Scale bars, 10 μm.

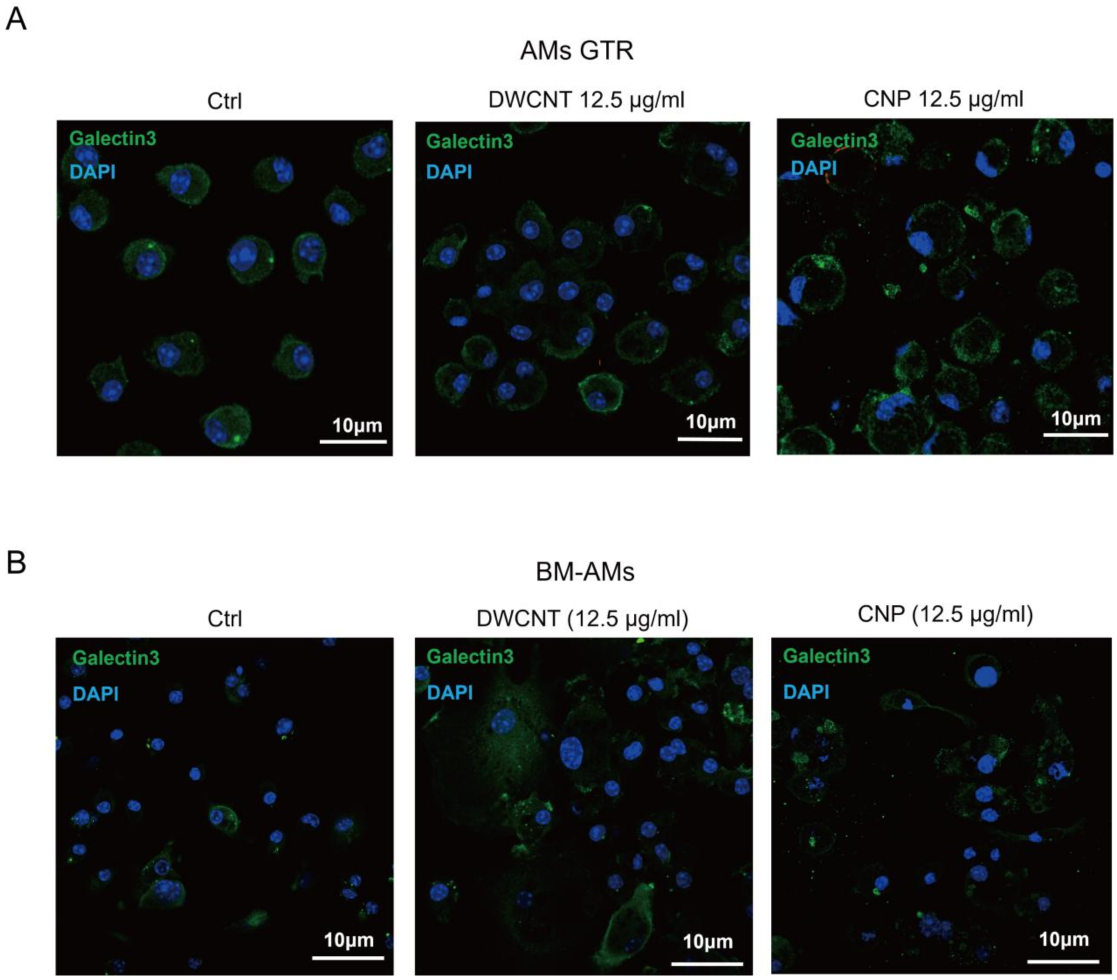


**Supplementary Fig. S6. Galectin-3 staining following CBNs exposure in AMs GTR and BM-AMs**

**(A, B)** Representative immunofluorescence images of Galectin-3 staining in AMs GTR and BM-AMs, respectively, under control conditions or following exposure to DWCNT (12.5 μg/ml) or CNP (12.5 μg/ml). Galectin-3 is shown in green, and nuclei were counterstained with DAPI (blue). Scale bars, 10 μm.

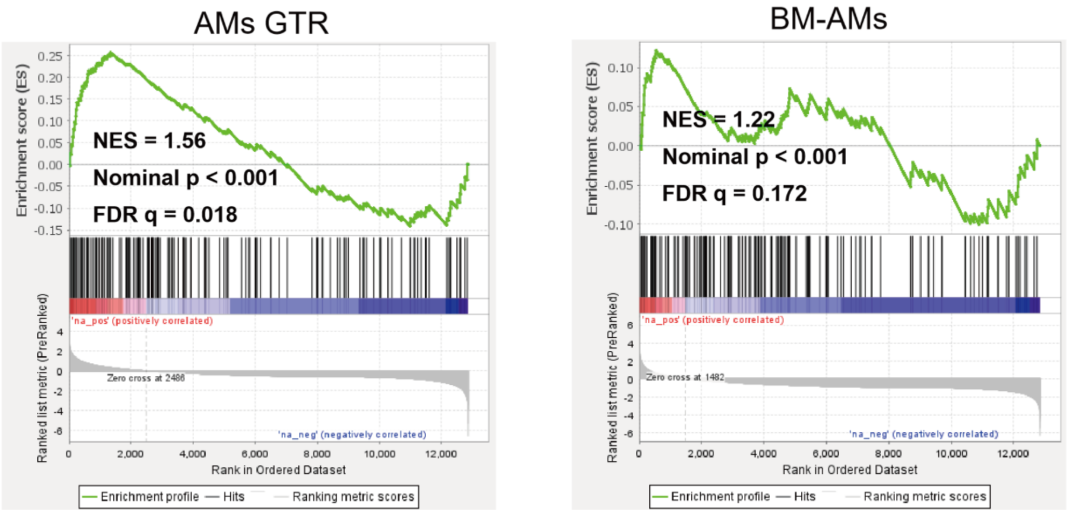


**Supplementary Fig. S7. Glycolysis-related gene set enrichment in AMs GTR and BM-AMs relative to in vivo alveolar macrophages**

Gene set enrichment analysis (GSEA) of the Hallmark Glycolysis gene set in AMs GTR and BM-AMs compared with scRNA-seq-derived *in vivo* AMs. AMs GTR showed moderate positive enrichment of glycolysis-related genes (**NES = 1.56, nominal P < 0.001, FDR q = 0.018**), whereas BM-AMs showed a more modest positive enrichment (**NES = 1.22, nominal P < 0.001, FDR q = 0.172**). Positive NES values indicate enrichment toward the *in vitro* AM models relative to *in vivo* AMs.
